# Adolescent frontal top-down neurons receive heightened local drive to establish adult attentional behavior in mice

**DOI:** 10.1101/2020.05.26.117929

**Authors:** Elisa M Nabel, Yury Garkun, Hiroyuki Koike, Masato Sadahiro, Ana Liang, Kevin J Norman, Giulia Taccheri, Michael P. Demars, Susanna Im, Keaven Caro, Sarah Lopez, Julia Bateh, Patrick R. Hof, Roger L. Clem, Hirofumi Morishita

## Abstract

Frontal top-down cortical neurons projecting to sensory cortical regions are well-positioned to integrate long-range inputs with local circuitry in frontal cortex to implement top-down attentional control of sensory regions. How adolescence contributes to the maturation of top-down neurons and associated local/long-range input balance, and the establishment of attentional control is poorly understood. Here we combine projection-specific electrophysiological and rabies-mediated input mapping in mice to uncover adolescence as a developmental stage when frontal top-down neurons projecting from the anterior cingulate to visual cortex are highly functionally integrated into local excitatory circuitry and have heightened activity compared to adulthood. Chemogenetic suppression of top-down neuron activity selectively during adolescence, but not later periods, produces long-lasting visual attentional behavior deficits, and results in excessive loss of local excitatory inputs in adulthood. Our study reveals an adolescent sensitive period when top-down neurons integrate local circuits with long-range connectivity to produce attentional behavior.

## Main Text

### Introduction

Altricial species, including both humans and mice, are born with relatively immature brains that follow a postnatal developmental trajectory before adult cognitive capacities are established. During development, initial axonal and dendritic branching is followed by a stage of synaptic overproduction and reduction to an adult plateau level ^1^. Particularly in the frontal cortex, this postnatal maturational process is integral for corticocortical projection neurons to integrate local inputs with long-range distal inputs carrying sensory, motor, and cognitive information and consequently to execute essential complex behaviors. Notably, deficits in frontal cortex long-range connectivity are pervasively reported in neurodevelopmental disorders and psychiatric disorders with developmental origins^2-4^, and often emerge following childhood and adolescence ^5-8^. Yet limited knowledge of normal processes governing corticocortical circuit development hinders further pathophysiological insight into these conditions.

Of particular significance among corticocortical projections is the top-down feedback projection from frontal cortex to sensory visual cortex^9,10^. Non-human primate studies demonstrate that a frontal projection to visual cortical areas executes attention controlled downstream visual processing ^9^. In mice, the anterior cingulate cortex area (ACA) and adjacent secondary motor cortex area (MOs) of the frontal cortex has been shown to project to visual cortical areas including primary visual cortex (VIS) ^11-13^. Optogenetic terminal stimulation of ACA neurons projecting to VIS (ACAVIS) in VIS can enhance the signal to noise ratio of neural response to preferred visual orientation stimuli and therefore increase the visual cortex ability to detect visual features—a hallmark feature of visual attention ^11^. Of note, disruption of functional connectivity between frontal cortex and visual cortex during visual attention tasks is implicated in several neurodevelopmental disorders, including schizophrenia, autism spectrum disorder, and attention deficit hyperactivity disorder ^2,14-16^. Here we aim to identify the circuit basis of adolescent maturation of mouse top-down ACA_VIS_ projection neurons and interrogate the contribution of this developmental window to adult circuitry and cognitive behavior.

Among various cognitive behaviors, we focus on visual attentional behavior by utilizing the five-choice serial reaction time task (5CSRTT), which is known to require ACA activity ^17^, in order to dissect the functionality of ACA_VIS_ in the context of visual attention. Adapted from the human continuous performance attention task in humans, the 5CSRTT is extensively used to examine visual attention in rodents ^18-22^. The task requires the subject to touch one of five spatial locations in a horizontal array of apertures on touchscreen when it flashes in order to receive a reward. Because the stimulus is presented pseudorandomly across these five spatial locations, the subjects must divide their attention across the areas. The number of times the subject touches the screen that flashed, in relationship to missed and incorrect touches, reflects a level of attentional processing^23^. Here, we integrate electrophysiology, input mapping, chemogenetics, and a translational visual attentional behavioral task in circuit-specific manner in mice, and demonstrate an adolescent sensitive period when top-down cortical circuits refine their local excitatory inputs in an activity-dependent manner to establish adult visual attention behavior.

## Results

### Greater adolescent excitatory drive onto top-down neurons

We first characterized the localization of top-down projection neurons projecting from frontal cortex to VIS in the adult mice. Retrograde viral labeling of these projection neurons to VIS revealed that a majority of labeled medial frontal neurons arise from dorsal ACA (ACAd) (65%), followed by adjacent secondary motor area (MOs) (27%) **(Supplementary Figure 1 a-c)**. Quantification of cortical layer distribution of these neurons showed approximately 81% arising from layers V/VI and 19% arising from layers II/III in adulthood **(Supplementary Figure 1 d)**. In the following we thus focused on characterizing top-down projection neurons from ACA to VIS (ACA_VIS_ projection neurons).

We next assessed changes in synaptic drive onto top-down ACA_VIS_ projection neurons during postnatal development across adolescent age. Adolescence is characterized by various behavioral and physiological changes that differs substantially from both infancy and adulthood, and broadly correspond to a period after the completion of weaning (p22-25) before the beginning of adulthood (p55-65)^24,25^. Of note, adolescent period around p32 is characterized by an increase in social play and affiliative behaviors ^26^. We performed whole-cell patch clamp recordings of miniature excitatory (mEPSC) and inhibitory (mIPSC) postsynaptic currents from ACA_VIS_ projection neurons in juvenile (∼p20: p18-p22), adolescent (∼p32: p29-p37), and adult (∼p78: p65-p100) mice. Fluorescent retrobeads were injected into the VIS 4-6 days prior to slice experiments to visualize ACA_VIS_ projection neurons **(Figure 1a)**. While there was no significant change in mEPSC frequency between the juvenile to adolescent period, we observed a significant decrease in mEPSC frequency from adolescence to adulthood **(Figure 1b and d)**, suggesting adolescence as a period of heightened excitatory drive. The cumulative distribution of mEPSC amplitudes shifted toward higher values between juvenile and adolescent groups with no changes in distribution between adolescents and adults, and mean amplitude of mEPSC also did not differ between the adolescent period and adulthood **(Figure 1b and e**). mIPSC frequency showed no statistically significant changes across ages **(Figure 1c and f**). Similar to mEPSC, the cumulative distribution of mIPSC amplitudes shifted toward higher values between juvenile and adolescent stages with no changes in distribution between adolescents and adults, and mean amplitude of mIPSC did not differ among three groups **(Figure 1e and g)**. Overall, our data show a selective shift in the frequency of excitatory postsynaptic currents between the adolescent and adult groups with no changes in amplitude after adolescence. In contrast, mIPSC frequency and amplitude showed no significant changes across age groups. Collectively, our study highlights adolescence as a period of greater adolescent excitatory drive onto frontal top-down ACA_VIS_ projection neurons.

**Figure 1:**
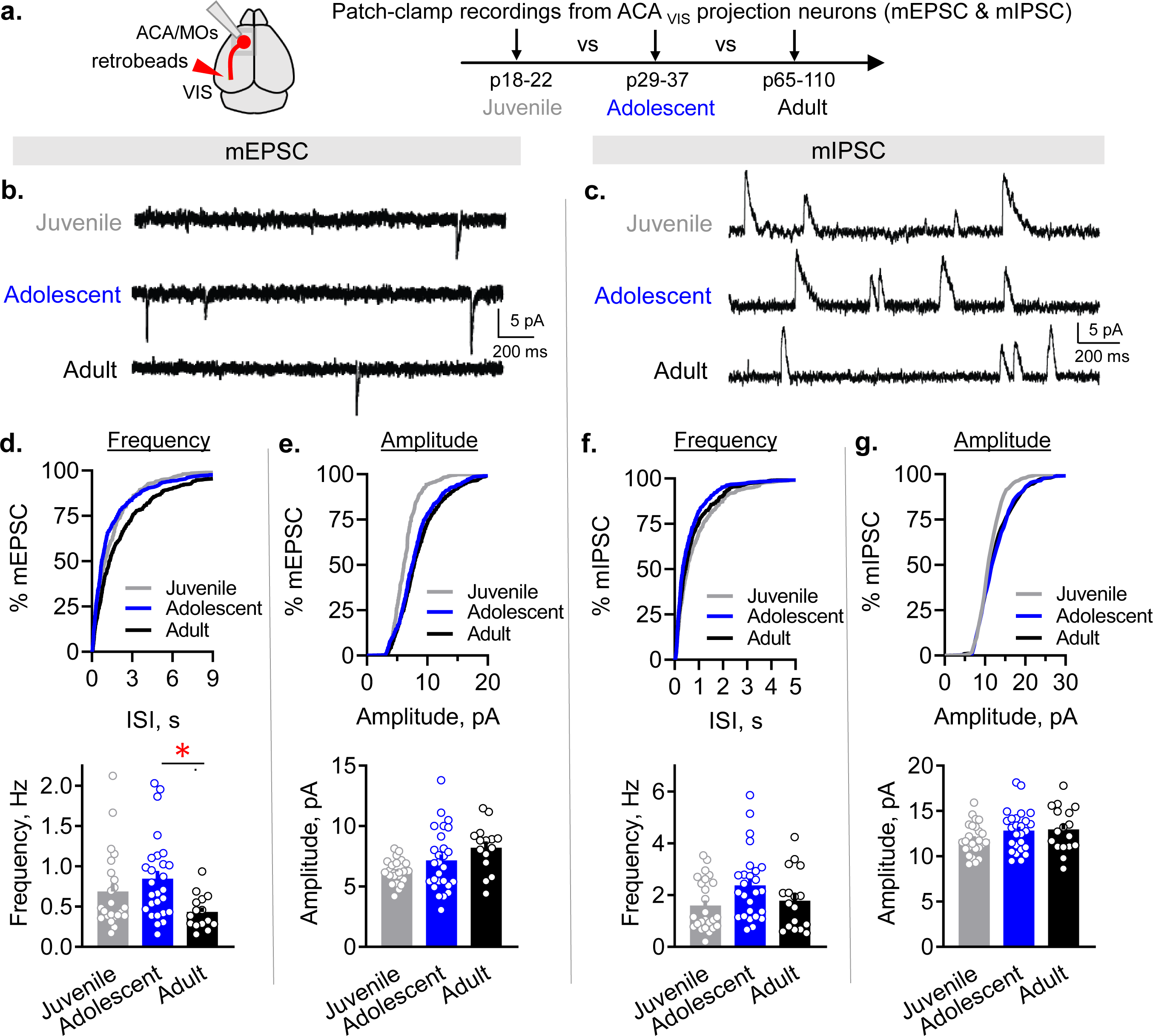
Adolescent ACA_VIS_ projection neurons receive greater excitatory drive. **a**. Developmental changes in excitatory and inhibitory synaptic drive onto top-down ACA_VIS_ projection neurons were assessed cross-sectionally. Slice patch-clamp recordings were performed from ACA_VIS_ projection neurons, fluorescently labeled by retrobeads injected in the VIS, on ACA/MOs brain slices at p20.3±-0.3 in average (p18-22) for Juvenile group, at p32.7±0.4 in average (p29-37) for Adolescent group, and at p78.1±4.3 in average (p65-110) for Adult group. Miniature excitatory postsynaptic currents (mEPSC) and miniature inhibitory postsynaptic currents (mIPSC) were recorded in the presence of TTX at −70 mV and 0 mV respectively. **b-c**. Representative traces of mEPSC (**b**) and mIPSC (**c**) measured from ACA_VIS_ neurons in three developmental age groups (for mEPSC: Juvenile: n=23 cells from 5 biologically independent mice, Adolescent: n=27 cells from 11 biologically independent mice, Adult: n=15 cells from 6 biologically independent mice; for mIPSC: Juvenile: n=25 cells from 5 biologically independent mice, Adolescent: n=26 cells from biologically independent 11 mice, Adult: n=17 cells from 6 biologically independent mice). Cumulative distribution (top) and averaged data (bottom) of mEPSC frequency (**d)** and mEPSC amplitude (**e**) in three age groups. Average mEPSC frequency were significantly higher in adolescent mice compared to adult (one-way ANOVA *F*_2,62_=4.099, *P*=0.021, post hoc Tukey test *P*_Juv-Adoles_ =0.43, \**P*_Adoles-Adult_ =0.016). While average mEPSC amplitude shows overall age effect, there was no significant difference between adolescence and adulthood (one-way ANOVA *F*_2,62_=4.573, *P*=0.014; post hoc Tukey test *P*_Juv-Adoles_=0.24, *P*_Adoles-Adult_=0.23). Cumulative distribution (top) and averaged data (bottom) of mIPSC frequency (**f)** and mEPSC amplitude (**g**) in three age groups show trending yet no significant difference in average mIPSC frequency (one-way ANOVA *F*_2,65_=3.060, *P*=0.054), and no difference in average mIPSC amplitude (one-way ANOVA *F*_2,65_=1.773, *P*=0.18). Data in d-g are presented as mean +/- s.e.m. See related Supplementary Figure 1. Source data are provided as a Source Data file.

### Heightened local inputs onto adolescent top-down neurons

To characterize this shift in excitatory drive between adolescence and adulthood further, we next sought to identify the input source onto frontal top-down ACA_VIS_ neurons present during adolescence that is absent by adulthood. We employed rabies mediated monosynaptic input mapping that would allow us to detect brain regions providing inputs to frontal ACA_VIS_ projection neurons and to quantify if neural input from these brain regions changed across development. We used a pseudotyped and genetically modified rabies virus (RbVdG(EnvA)-eGFP) that allows visualization of starter cells by requiring the cognate receptor, avian tumor virus receptor A (TVA), to be expressed for cell entry and rabies glycoprotein (RbG) for cell exit ^27^. Specifically, this system restricts expression of GFP only in starter ACA_VIS_ projection neurons expressing TVA and their presynaptic inputs. ACA_VIS_ circuit specific TVA and RbG expression was conferred by an intersectional viral approach that introduced Cre-dependent TVA and RbG vectors into the ACA and retrograde canine adenovirus type 2 (CAV-2) Cre in VIS. This approach targeted ACA_VIS_ neurons mainly in deep layers consistent with the characteristic of CAV-2 transduction pattern^28^, but also some in superficial layers comparable to the labeling pattern of retrobeads or retro AAV2 **(Figure 2a)**. After allowing 2.5 weeks for gene expression, monosynaptic input labeling was accomplished by injection of RbVdG(EnvA)-eGFP. We examined expression 7 days later and estimate that monosynaptic input labeling at adolescent and adult time points represent connectivity around ages p34 and p80 respectively **(Figure 2b)**. Because the p12 and p60 brain differed in size when stereotactic surgery was performed, we first characterized and compared the distribution of starter cells initiating the retrograde transfer of rabies virus, identified as double-labelled mCherry+ eGFP+ **(Figure 2c)** between adolescent and adult groups and found no difference in their anterior-posterior distribution **(Figure 2d)**, or in the absolute number of starter cells per animal across the two age groups **(Figure 2e)**. An additional experiment verified that the distribution of ACA_VIS_ neurons across the cortical layers does not change across the age groups **(Supplementary Figure 1d)**. Consequently, we were able to directly compare the number of monosynaptic input neurons from each anatomical location between adolescent and adult groups by normalizing to the number of starter cells for each animal.

**Figure 2:**
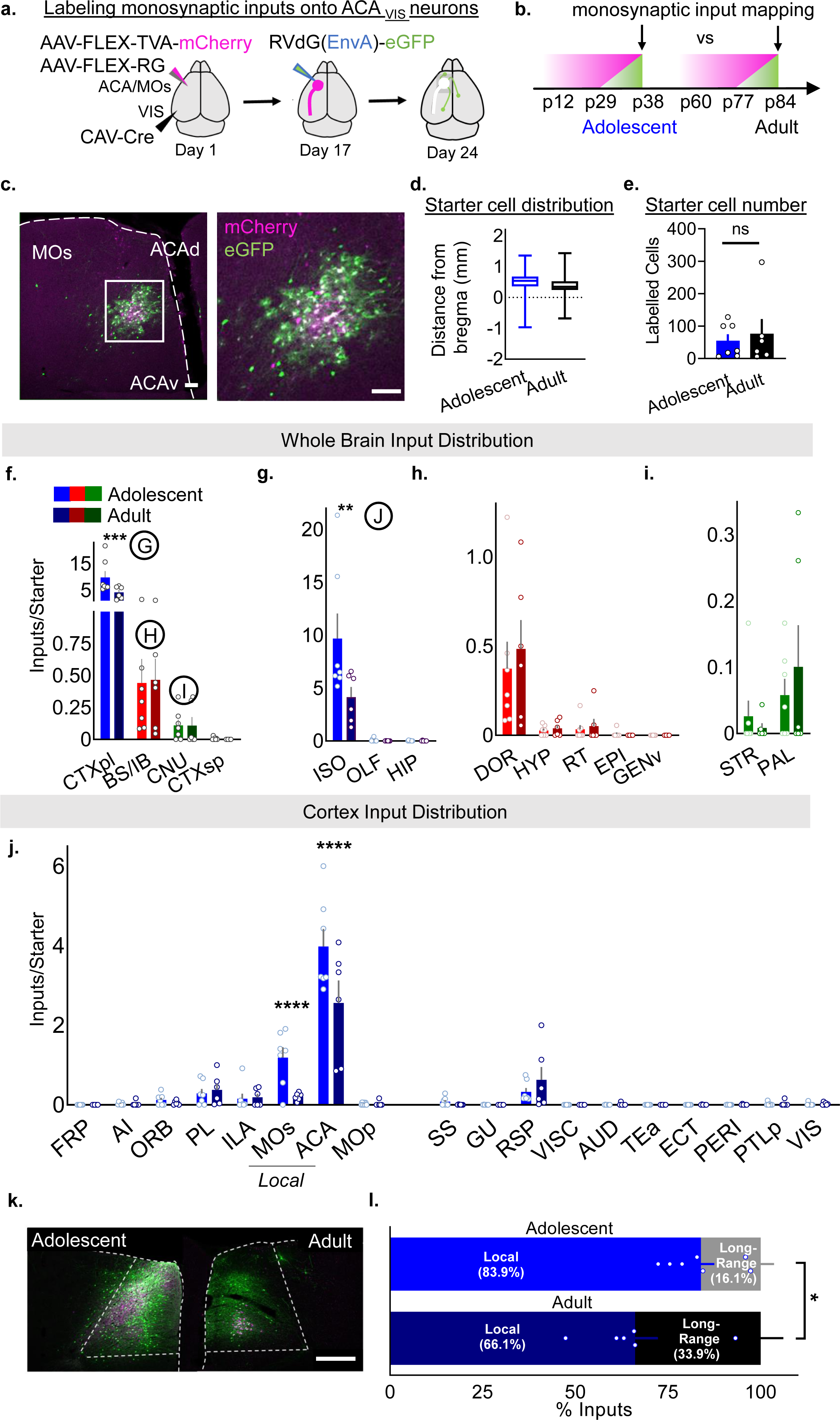
Adolescent ACA_VIS_ neurons receive greater local presynaptic inputs. **a**. Schematic overview of intersectional viral injection approach enabling labeling of presynaptic input neurons specifically onto top-down ACA_VIS_ projection neurons. Cre-dependent AAVs expressing TVA-mcherry and Rb-G were injected into the ACA and retrograde CAV-Cre into VIS. After 2.5 weeks, pseudotyped rabies virus was injected into the ACA to allow for uptake of rabies virus in top-down neurons only to express GFP over one week. **b**. Injection timeline for two age groups. AAV injections were initiated at p12 to capture inputs during adolescence (p38) and at p60 to capture inputs during adulthood (p84). **c**. Top-down starter cells and local inputs were visualized in ACA. Left: Magenta helper TVA-mcherry infected cells permit rabies entry. White starter AAV TVA-mcherry + Rbs-GFP co-infected cells allow rabies to retrograde back one synapse onto green monosynaptic input cells. Right: High magnification of boxed area. Scale Bars =100 µm. Experimental images were obtained from 13 mice, with similar results obtained. **d**. The anterior posterior distribution of starter cells is consistent across adolescent (n=7 biologically independent mice) and adult groups (n=6 biologically independent mice). Box plot shows the median, the interquartile range, and whiskers (minima to maxima). (nested two-tailed unpaired *t*-test; *t*_11_=0.167, *P*=0.988, n= 386 cells from 7 biologically independent mice [adolescent group], 474 cells from 6 biologically independent mice [adult group]). **e**. The absolute number of starter cells per animal are comparable across the two age groups (two-sided unpaired *t*-test, *t*_11_=0.4634, *P*=0.6521, n=7 [adolescent], 6 [adult] biologically independent mice). **f-j**. Absolute total number of input neurons in adolescence (n=6 biologically independent mice) and adulthood (n=7 biologically independent mice) normalized to starter cell count per animal. See **Supplementary Table 1** for abbreviations. **f**. Compared to adult mice, adolescent mice show a larger number of input cells at the highest hierarchical level of neuroanatomical organization selectively in the cortical plate (CTXpl) (two-way RM ANOVA, age x brain region interaction *F*_3,33_=4.091, *P*=0.0142 post hoc analysis Sidak’s multiple comparisons test \*\*\**P*=0.0008). No inputs were detected from the midbrain or hindbrain for either group. **g**. Within the CTXpl, adolescent mice showed a larger number of input cells selectivity in the isocortex (ISO) (two-way RM ANOVA, age x brain region interaction *F*_2,22_=4.168, *P*=0.0292, post hoc analysis Sidak’s multiple comparisons test, \*\**P*=0.0036), and not in the olfactory area (post hoc analysis, *P*=0.999) or hippocampus (post hoc analysis, *P*=0.999). **h**. Adolescent and adult mice show the same number of inputs coming from sub-regions of the diencephalon (two-way RM ANOVA, age x brain region interaction *F*_4,44_=0.2566, *P*=0.9041), and **i**. cerebral nuclei (two-way RM ANOVA, age x brain region interaction *F*_1,11_=1.097, *P*=0.3173). **j**. In the isocortex, more input neurons are found selectively in the secondary motor area (MOs) (two-way RM ANOVA, age x brain region interaction *F*_17,187_=4.122, *P*=0.524 x10^−6^, post hoc analysis Sidak’s multiple comparison test \*\*\*\**P*=0.532 x10^−4^) and anterior cingulate cortex (ACA) (post hoc analysis \*\*\*\**P*=0.841 x10^−9^). **k**. Representative image of ACA and MOs in adolescent vs adult group (Scale bar 500µm). Experimental images were obtained from 7 or 6 mice respectively, with similar results obtained. **l**. Local:Long-Range input ratio is higher in adolescence compared to adults (unpaired *t*-test, *t*_11_=2.586, \**P*=0.0253, n=7,6 biologically independent mice). Data in e-j, l are presented as mean +/- s.e.m. See related Supplementary Figure 2. Source data are provided as a Source Data file.

In each animal, all monosynaptic input neurons for evenly sampled coronal slices were counted and assigned to the corresponding brain regions of the Allen Brain Atlas hierarchical neuroanatomic ontology (http://mouse.brain-map.org/static/atlas) **(Supplementary Table1)**. To allow a direct comparison of input number and distribution across animals and developmental groups, for each animal, inputs were normalized by dividing by the number of starter cells. At the highest hierarchical level of organization, we observed more inputs in the cortical plate (CTXpl) in the adolescent group with no differences observed in the brainstem, where all inputs were limited to the diencephalon (BS/IB), the cerebral nuclei (CNU), and the cortical subplate (CTXsp). No inputs from the midbrain (MB) or hindbrain (HB) were detected **(Figure 2f)**. At the next nested level of brain regions, we found that changes in the CTXpl were contained entirely within the isocortex (ISO) **(Figure 2g)**. No developmental differences within the BS/IB and CNU were found **(Figure 2h and 2i)**. Within the ISO, only two brain regions, the ACA and anatomically adjacent secondary motor cortex (MOs) showed a significant difference between the adolescent and adult groups **(Figure 2j)**. Specifically, the ACA and MOs provided more inputs at adolescence, suggesting that these local inputs are reduced by adulthood **(Figure 2k)**. Although this system is noted to have local, non-RbG-dependent labeling ^29^, we found that the amount of labeling in the presence of RbG **(Supplementary Figure 2a)** was much greater than an independent experiment in which RbG was excluded, and that there was no difference in leak between adolescent and adult groups **(Supplementary Figure 2b)**. This was additionally confirmed by an independent experiment in which RVdG(EnvA)-eGFP was injected in VIS for uptake by TVA expressed at axon terminals to avoid local leak **(Supplementary Figure 2c)**.

As these data revealed a reduction of input source following adolescence, we next set out to determine the impact of this structural switch on top-down ACA_VIS_ projection integration into its neural networks. We calculated the relative percentage of total inputs coming from local ACA and MOs areas compared to all other long-range distal brain regions for each animal. Our analysis revealed that the relative ratio of local to long-range distal inputs is significantly higher in adolescents (5.21:1) compared to adults (1.74:1) **(Figure 2l)**. Together these results support a model that top-down ACA_VIS_ neurons receive greater local inputs during adolescence than adulthood, which in turn increase a relative contribution of long-range distal connectivity into adulthood.

### Greater local excitatory synaptic inputs to top-down neurons

Our patch-clamp recordings (**Figure 1**) and rabies mediated mapping of monosynaptic input onto ACA_VIS_ projection neurons (**Figure 2**) collectively point to the model that ACA_VIS_ neurons are more extensively connected with local frontal excitatory networks during adolescence than adulthood. As rabies mapping cannot fully rule out non-specific labeling within the local injection area ^29^ as well as an activity-dependent labelling ^30^, we next set out to perform non-rabies based approach to directly examine local functional connectivity. To this end, we combined optogenetics and slice patch-clamp recording by applying Cre- and Flippase (Flp)-dependent intersectional viral strategy called INTRSECT ^31^ to allow optogenetic interrogation of local inputs selectively without direct activation of ACA_VIS_ projection neurons (**Figure 3a**). This approach can overcome the challenge of assessing the very low probability of connectivity between pairs of ACA pyramidal neurons (2% in recent study ^32^) using conventional paired patch clamp recordings. We infused retrograde Cre encoding virus in VIS to retrogradely label the soma of ACA_VIS_ projection neurons in ACA, and also introduced codon-optimized Flp (Flpo) and FlpON/CreOff-ChR2 vectors ^31^ into the ACA to express ChR2 selectively in Flpo-expressing ACA neurons yet actively avoid labeling ACA_VIS_ projection neurons expressing Cre. AAV infusion was performed around p12 or p43 for adolescence and adult recordings respectively. After waiting ∼18 days for ChR2 accumulation, we performed whole-cell patch-clamp recordings. The spread and intensity of YFP signal of analyzed slices show comparable levels of ChR2 expression between groups (**Figure 3b, c**; see details in methods section). Patch-clamp recordings were obtained from visually identified mCherry+ACA_VIS_ neurons in acute ACA slices obtained from adolescent and adult mice. Brief pulses of blue light (0.5 ms) successfully activated local inputs onto ACA_VIS_ neurons and elicited reliable EPSCs in every ACA_VIS_ neuron recorded. The latency from light onset to the onset of EPSCs in ACA_VIS_ neurons was 5.2 ms on average due to synaptic delay, suggesting monosynaptic connectivity. In our experiments, we found no cells with the latency below 2 ms which are considered ChR2-expressing.

**Figure 3:**
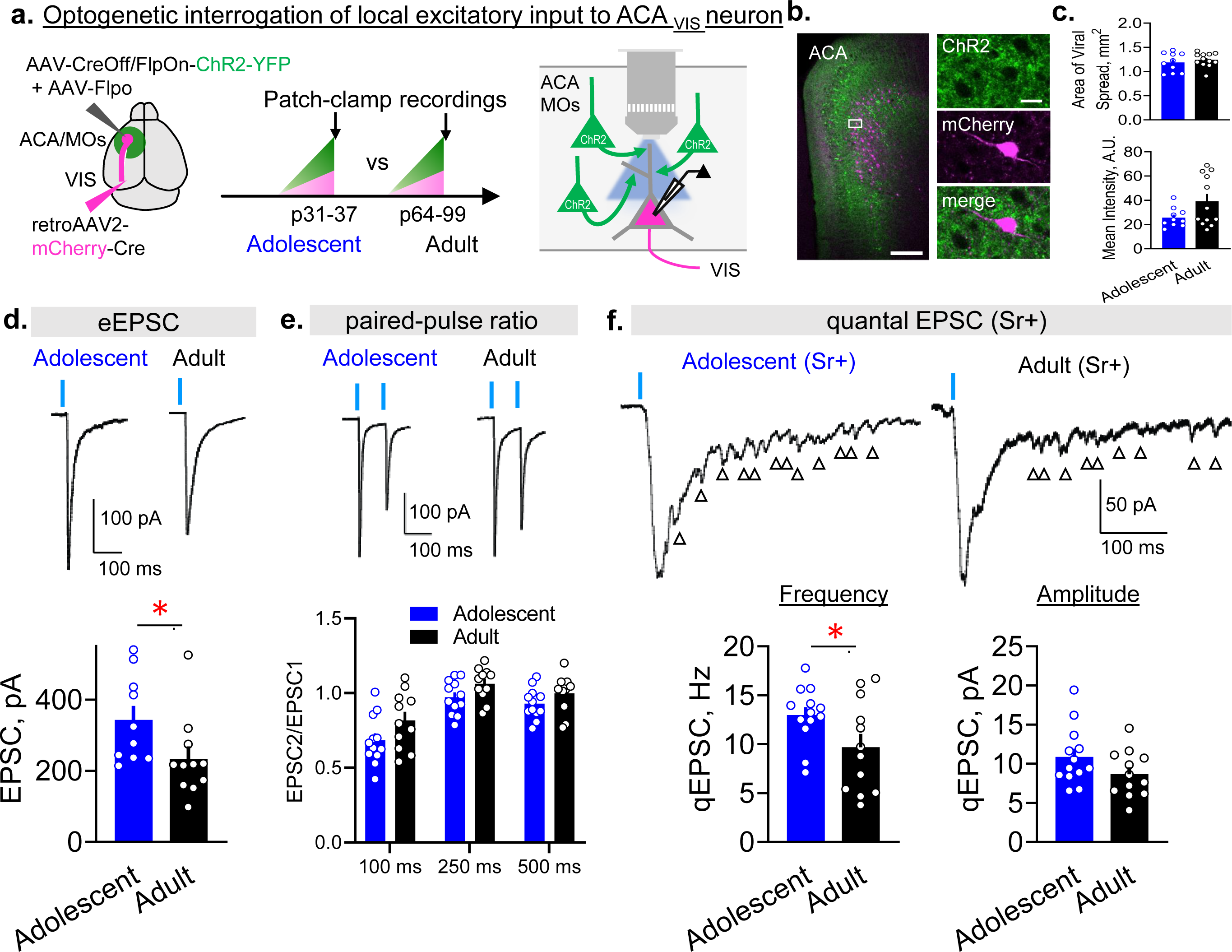
Greater adolescent local excitatory synaptic inputs onto ACA_VIS_ neurons. **a**. Schematic of intersectional viral approach to test local excitatory synaptic input specifically onto ACA_VIS_ projection neurons in adolescence and adulthood. (left) Injections of retrograde AAV2-mCherry-Cre into the VIS labeled ACA_VIS_ projection neurons in ACA with red fluorescence, while injections of AAV-Flpo and AAV-CreOff/FlpOn-ChR2- YFP into ACA enable ChR2-YFP expression in all but ACA_VIS_ projection neurons. (middle) AAV injections were done at p12.8±0.5 in average (p11-15) for Adolescent group and at p43.0±1.9 in average (p38-58) for Adult group. Slice patch clamp recordings were performed at p33.5±0.7 in average (p31-37) for Adolescent group and at p71.8±4.2 in average (p64-99) for Adult group. (right) Local inputs were stimulated in situ by activating ChR2 using wide-field LED. **b**. (left) Labeling of local inputs (green) and ACA_VIS_ projection neurons (magenta) in ACA. (right) High magnification of boxed area showing mCherry+ ACA_VIS_ neuron does not express ChR2-YFP and is surrounded by ChR2+YFP terminals. Scale bars: 200 μm on the left and 20 μm on the right. Experimental images were obtained from 17 mice with similar results obtained. **c**. Quantification of the spread and intensity of the YFP signal shows no significant differences between Adolescent and Adult groups (area: two-tailed unpaired *t*-test, *t*_20_=0.756, *P*=0.4585, mean intensity: two-tailed unpaired *t*-test, *t*_20_=1.936, *P*=0.0672, n=11 slices from 10 biologically independent mice for adolescent and n=12 slices from 7 biologically independent mice for adult groups). **d**. Light-evoked monosynaptic EPSC traces (top) and mean amplitude (bottom) for two developmental age groups (two-tailed unpaired *t*-test, *t*_19_=2.122, \**P*=0.0472, n=10 cells from 5 biologically independent mice for adolescent and n=11 cells from 6 biologically independent mice for adult groups). Blue dashes indicate the time of light stimulation. **e**. Short-term dynamics of local synaptic inputs onto ACA_VIS_ neurons. Representative EPSC evoked by pair of light stimuli (top) and average paired pulse ratio (bottom) (EPSC2/EPSC1 Adolescent vs. EPSC2/EPSC1 Adult, two-way ANOVA, time x age interaction: *F*_*2*,63_=0.3418, *P*=0.7118, n=12 cells from 4 biologically independent mice for adolescent and n=11 cells from 4 biologically independent mice for adult). **f**. Examples of light-evoked quantal EPSC (qEPSC) in ACA_VIS_ neurons in the presence of strontium (top) to produce delayed asynchronous quantal release of neurotransmitters. Arrows indicate detected qEPSCs. (bottom) Averaged qEPSC amplitude and frequency for Adolescent and Adult groups (two-tailed unpaired *t*-test, frequency: *t*_23_=2.118, \**P*= 0.0452, amplitude: *t*_23_=1.626, *P*=0.12, n=13 cells from 4 biologically independent mice for adolescent and n=12 cells from 5 biologically independent mice for adult). Data in c-f are presented as mean +/- s.e.m. Source data are provided as a Source Data file.

First, to assess evoked excitatory monosynaptic transmission from local inputs, we isolated monosynaptic connections with tetrodotoxin (TTX, 1μM), 4-AP (100 μM) and picrotoxin (100 μM). Light intensity was adjusted to elicit stable response that didn’t change with the additional increase of the stimulus intensity. We found that the amplitude of evoked EPSC was higher in adolescent animals in comparison to the adults (**Figure 3d**). Differences in the amplitude of evoked responses could be due to changes in presynaptic release probability, changes in synaptic strength or synapse number. To investigate more thoroughly, we next used paired stimulation to evoke EPSCs and quantify possible changes in short-term dynamics. Experiments were performed in the absence of TTX, 4-AP and picrotoxin to preserve presynaptic action potentials. In response to paired stimulation, we observed short-term depression of local inputs to ACA_VIS_ neurons (**Figure 3e**). In agreement with the published data ^33^, magnitude of short-term depression depended on the interstimulus interval with the higher depression at shorter intervals. Paired-pulse analysis did not show significant interaction between age and inter-pulse interval (**Figure 3e**), suggesting that the change in EPSCs could be due to changes in synaptic strength or synapse number. To further differentiate between these possibilities, we recorded EPSCs in ACSF where extracellular calcium (Ca^2+^) was replaced with strontium (Sr^2+^, 3 mM) in the presence of TTX, 4-AP and picrotoxin (**Figure 3f**). Sr^2+^ desynchronizes glutamate release and allows the resolution of quantal synaptic events. The resultant local input-specific qEPSC provides a measure of unitary synaptic strength (amplitude) and a number of connections (frequency) ^34 35^ Optogenetic stimulation of local ACA excitatory axons revealed higher amount of quantal synaptic events in adolescent animals in comparison to adult (**Figure 3f**). In contrast, we found no difference in the amplitude of light-evoked quantal events (**Figure 3f**). Collectively, these results suggest that, during adolescence, the local excitatory drive onto ACA_VIS_ neurons is higher in adolescence than in adulthood likely due to the increased synaptic number from local excitatory inputs.

### Heightened adolescent *in vivo* activity of top-down neurons

As ACA_VIS_ neurons are more extensively connected with local frontal excitatory networks during adolescence than adulthood, we next investigated if ACA_VIS_ projection neurons themselves have heightened activity during adolescence compared to adulthood **(Figure 4a)**. To evaluate ACA_VIS_ neuron activity *in vivo*, we expressed channelrhodopsin-2 (ChR2) as an optogenetic tag to allow efficient identification of individual ACA_VIS_ neurons amongst a population of neurons by using a blue light (473 nm) as a search stimulus during *in vivo* electrophysiological recordings **(Figure 4c)**. We targeted expression of ChR2-GFP in ACA_VIS_ projection neurons by injecting retrograde AAV2-ChR2-GFP into the VIS **(Figure 4a, b)**. After >2 weeks of viral incorporation, the mice were subjected to *In vivo* electrophysiological recordings at adolescent (∼p36: p33-39) and adulthood (∼p79: p76-82). Using this technique, detected units were confirmed as ACA_VIS_ neurons if they also responded to optogenetic stimulation within 3 ms upon blue light stimulation **(Figure 4c-e)**. We found that there was a significantly greater spontaneous firing rate of ACA_VIS_ neurons *in vivo* during adolescence compared to adulthood **(Figure 4f,g)**. Non-tagged ChR2 (-) ACA neurons also showed heightened adolescent spiking activity compared to adulthood (**Supplementary Figure 3**), suggesting that greater adolescent activity may be a general feature of ACA neurons. However, it should be noted that we cannot rule out the possibility that some ACAvis neurons which do not express ChR2 are included in non-tagged ChR2 (-) ACA neuron population because our viral approach does not allow transduction of ChR2 in all ACAvis neurons. Taken together, our findings suggest that ACA_VIS_ projection neurons are not only highly functionally integrated into local excitatory circuitry within ACA **(Figure 1-3)**, but also ACA_VIS_ neurons themselves have heightened activity during adolescence.

**Figure 4:**
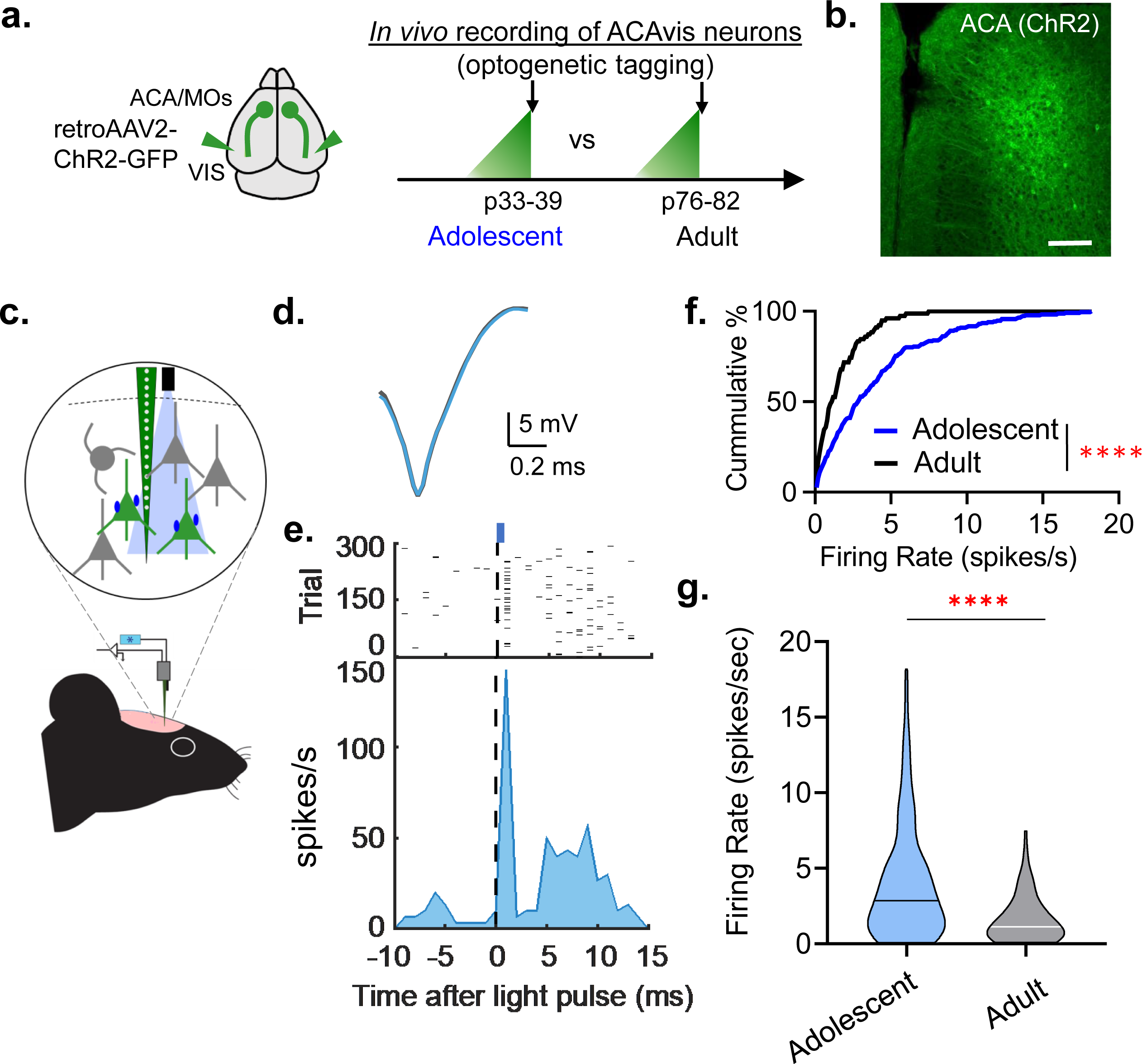
Greater firing activity of adolescent ACAvis neurons *in vivo*. **a**. Schematic of viral approach to express Channelrhodopsin-2 (ChR2) in frontal top-down ACAvis projection neurons for *in vivo* electrophysiological recordings of optically tagged ACAvis neurons in adolescent and adult mice. (left) Injections of retrograde AAV2-ChR2-GFP into the VIS enable ChR2-GFP expression in cells projecting to VIS including ACAvis projection neurons. (right) AAV injections were done at p20 for Adolescent group and at p63.3±0.7 in average (p62-64) for Adult group. Recordings were performed at p35.8±1.2 in average (p33-39) for Adolescent group and at p79.0±1.7 in average (p76-82) for Adult group. **b**. Representative images of viral GFP+ACAvis neurons in ACA following injection of retroAAV2-ChR2-GFP into the VIS. Scale: 200 µm. Experimental images were obtained from 8 mice, with similar results obtained. **c**. Schematic of *in vivo* electrophysiology setup with optical tagging. ChR2 has been used as an optogenetic tag to allow identification of ACAvis projection neurons (green) *in vivo* based on whether they can be activated by optical stimulation. **d**. Representative overlapping spike waveforms of optically-tagged ACAvis projection neuron recorded with (blue trace) and without (gray trace) optic stimulation. **e**. Raster plot and peristimulus time histogram showing time locked optogenetic activation of a representative ACAvis neuron within 2 ms. Laser (wavelength 473 nm, fiber diameter 105 μm) was delivered using an optic fiber coupled to the multichannel extracellular recording electrode. The fiber tip was positioned immediately above the ACA recording site. ACAvis neurons were identified with a light pulse (1 ms, 5Hz) stimulus. **f**. Cumulative distribution of spontaneous firing frequency of ACAvis projection neurons in adolescent and adult animals (blue line, n= 231 cells from 5 adolescent biologically independent mice, black line, n= 78 cells from 3 adult biologically independent mice, two-tailed Kolmogorov-Smirnov test, \*\*\*\**P*=0.141×10^−5^). **g**. Averaged firing activity of ACAvis projection neurons in adolescent and adult animals (blue, n=231 cells from 5 adolescent biologically independent mice, black, n= 78 cells from 3 adult biologically independent mice, two-tailed unpaired *t*-test, *t*_307_=5.379, \*\*\*\**P*=0.149×10^−6^). Horizontal lines within violin plots represent median values. See related Supplementary Figures 3. Source data are provided as a Source Data file.

### Adolescent top-down neuron activity promotes adult attention

We next examined whether this transient developmental stage of heightened ACA_VIS_ neuron activity had significance for adult function. Because ACA_VIS_ projection neurons can enhance the ability to detect visual features in VIS—a hallmark feature of visual attention ^11^, we aimed to determine if the activity of top-down ACA_VIS_ projection neurons during this adolescent period is required to establish adult visual attentional behavior. To this end, we perturbed ACA_VIS_ neuron activity during adolescence (p25-34) or late-adolescent (p45-54) periods by chemogenetically suppressing ACA_VIS_ neuron activity during these developmental windows and examined its long-lasting consequence on adult visual attentional behavior. Visual attention capacity was tested by the 5CSRTT^23^. A use of this unimodal visual attention task is particularly relevant for our study which focuses on the maturation of visual top-down ACA_VIS_ projection neurons, given that a previous chemogenetic silencing study demonstrated that 5CSRTT requires proper activity of ACA neurons in mice ^17^. In contrast, a cross-modal attention task, in which animals select between conflicting visual and auditory stimuli, does not require ACA or VIS activity ^36^. 5CSRTT assays sustain visual attention by requiring mice to maintain divided attention across 5 areas for the presentation of a brief solid while square visual stimulus presented randomly in one of 5 locations on black flat touchscreen that must be nose-poked to release a reward ^37^ **(Supplementary Figure 4)**. Prior studies have shown that attention deficits can be measured quantitatively as the combined error rate of an increase in omitted responses to light stimulus and a decrease in accurate responses ^20^. Our study was conducted in automated, standardized Bussey-Saksida touch-screen operant chambers that are highly analogous to Cambridge Neuropsychological Test Automated Battery for humans and non-human primates ^38-41^.

We introduced inhibitory designer receptor exclusively activated by a designer drug (iDREADD)^42^ into visual top-down ACA_VIS_ projection neurons **(Figure 5a)** using an intersectional Cre-dependent viral approach that introduced Cre-dependent DREADD vector into frontal cortex and retrograde Cre encoding virus in VIS (**Supplementary Figure 5**), and suppressed top-down neuron activity through Clozapine-N-oxide (CNO) delivered intraperitoneally twice daily during the adolescent period of heightened local connectivity from p25 through p34 **(Figure 5c)**. While our intersectional approach does not limit the expression of DREADD to frontal projection to primary sensory cortices due to collaterals to association sensory cortices ^43^, and therefore cannot rule out the contribution of these collaterals to behavior effects, a previous study demonstrated that stimulation of ACA_VIS_ projection terminals within primary sensory cortex (VIS) is sufficient to impact visual gain ^11^. To test the temporal specificity of this developmental perturbation, we also delivered CNO during the late-adolescent period (p45-p54) in a separate group of mice with top-down visual projection neuron iDREADD expression **(Figure 5c)**. As another control, we also examined the effect of suppressing an activity of an adolescent frontal population projecting to primary auditory cortex (AUD), which are located at secondary and primary motor cortical area (MOs/p) adjacent to ACAd ^43,44^ (MOs_AUD_ projection neurons) **(Figure 5 b, c)**. To confirm that neurons virally transfected at p12 were responsive to CNO during the adolescent period, we performed slice patch clamp recordings. Whole-cell current clamp recordings were obtained from mCherry expressing ACA_VIS_ neurons in ACA slices. A steady depolarizing current of 1.5-times rheobase (the minimal current to evoke action potential) was applied to initiate firing of ACA_VIS_ neurons following by application of CNO **(Supplementary Figure 6a)**. CNO application (10 μM) decreased the membrane potential **(Supplementary Figure 6b)** and inhibited spontaneous firing of projection neurons **(Supplementary Figure 6c)**.

**Figure 5:**
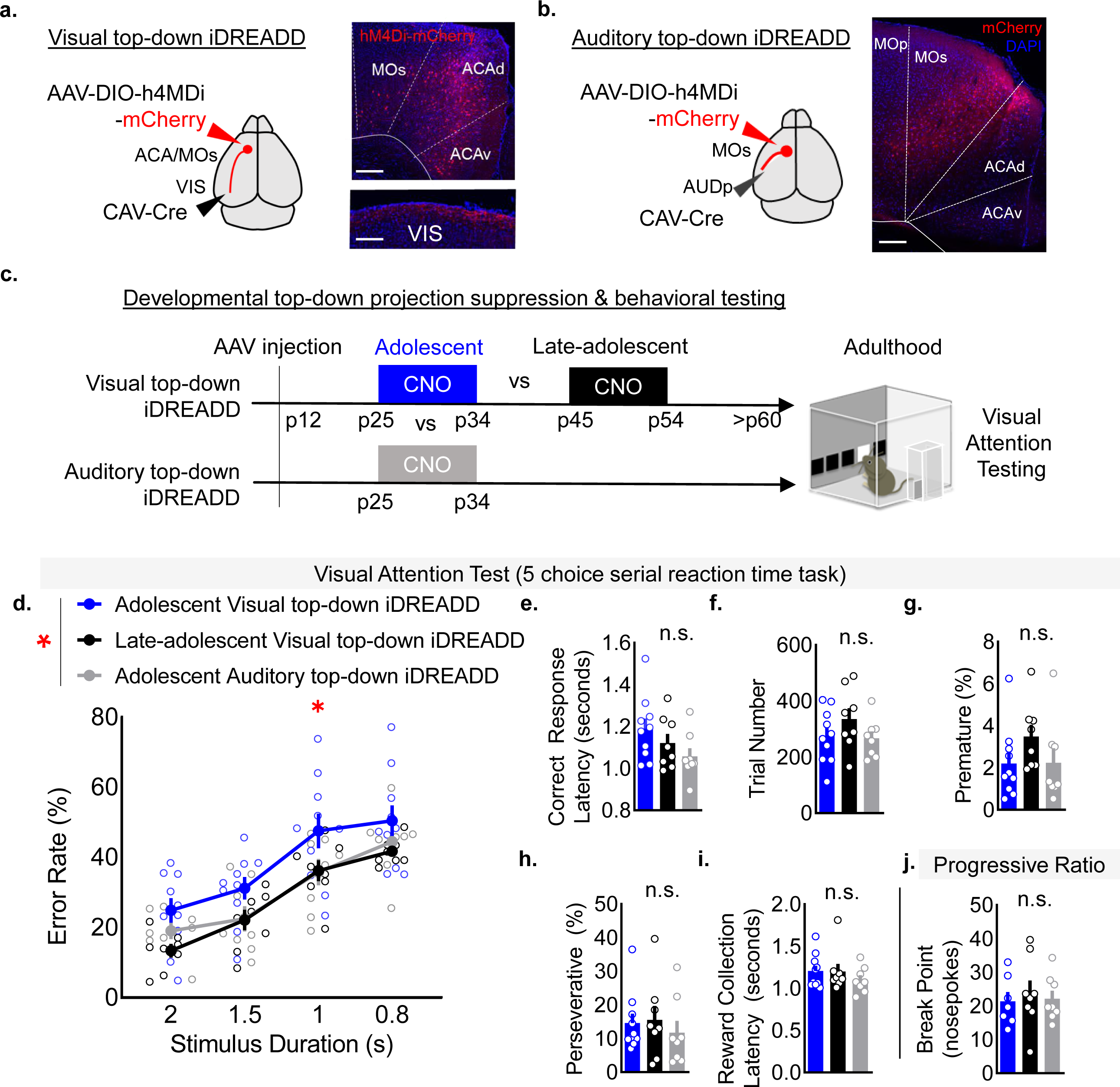
Suppression of adolescent ACA_VIS_ neuron activity produces a visual attention deficit in adulthood. **a-b**. (left) Schematic of intersectional viral approach to introduce hM4D(Gi)-inhibitory DREADD (iDREADD) selectively into **(a)** frontal visual top-down ACA_VIS_ projection neurons, or (**b**) frontal auditory top-down MOs_AUD_ projection neurons by introducing Cre-dependent iDREADD vectors into the frontal cortex and retrograde Cre encoding virus in VIS. Viruses were injected at p12 to allow for expression of iDREADD by p25. (right) Representative images of frontal cortex showing labeling of **(a**) mCherry+ACA_VIS_ projection neurons and their terminals in layer1 of VIS or (**b**) mCherry+MOs_AUD_ projection neurons in frontal cortex. Experimental images were obtained from 10 or 8 biologically independent mice respectively, with similar results obtained. Scale bars: 200µm. **c**. Overview of experimental timeline. Mice were injected with 10 mg/kg CNO in sterile saline twice a day from p25 to p34 or from p45 to p54 and started training on the five-choice serial reaction time task (5CSRTT) at adulthood (p60). See Supplementary Figure 2 for a detailed overview of the 5CSRTT. Mice were subsequently tested for motivation with the progressive ratio task. **d**. CNO-mediated suppression of frontal top-down projection neuron increases adult attentional error rate selectively in adolescent visual top-down suppression condition (two-way RM ANOVA, effect of group, *F*_2,23_ = 3.622, \**P*= 0.0429, Newman-Keuls multiple comparisons test: \**P*< 0.05: adolescent vs late-adolescent visual top-down suppression (n=10, 8 biologically independent mice), \**P*< 0.05: adolescent visual top-down suppression vs adolescent auditory top-down suppression (n=10,8 biologically independent mice)). **e-i**. Adolescent CNO administration does not alter other measures on the 5CSRTT in adult mice (n=10 [adolescent visual top-down iDREADD: blue bars], 8 [late-adolescent visual top-down iDREADD; black bars], 8 [adolescent auditory top-down iDREADD; gray bars] biologically independent mice): **e**. Correct response latency (one-way ANOVA, *F*_2,23_ = 2.204, *P*= 0.1331), **f**. Trial number (one-way ANOVA, *F*_2,23_ = 1.338, *P*= 0.2820) **g**. Premature responses (one-way ANOVA, *F*_2,23_ = 1.475, *P*= 0.2495) **h**. Perseverative responses (one-way ANOVA, *F*_2,23_ = 0.3161, *P*=0.7321) **i**. Reward collection latency (one-way ANOVA, *F*_2,23_ = 0.6789, *P*=0.5171) **j**. No differences in breakpoint were observed in the progressive ratio task (one-way ANOVA, *F*_2,20_ = 0.1693, *P*= 0.8454, n=7 [adolescent visual top-down iDREADD: blue bars], 8 [late-adolescent visual top-down iDREADD; black bars], 8 [adolescent auditory top-down iDREADD; gray bars] biologically independent mice). Data in d-j are presented as mean +/- s.e.m. See related Supplementary Figures 4-7. Source data are provided as a Source Data file.

During 5CSRTT testing, we observed a selective increase in error rate in mice with top-down ACA_VIS_ iDREADD that received CNO during adolescence compared to the group that received CNO during the late-adolescent period, or the MOs_AUD_ iDREADD group that received CNO during the same adolescent period **(Figure 5d)**. No differences in correct response latency **(Figure 5e)**, trial number **(Figure 5f)**, premature responses **(Figure 5g)**, or perseverative responses **(Figure 5h)** were observed, suggesting a selective visual attention deficit. An increase error rate did not reflect a decrease in motivation because the CNO group as they displayed no differences in latency to collect the reward during the task **(Figure 5i)**. Additionally, in an independent progressive ratio assay of motivation, no differences in break point to attempt release of the liquid reward **(Figure 5j)**. Further, no differences in acquisition of the 5CSRTT or baseline training performance prior to testing were observed **(Supplementary Figure 7)**. While it is challenging to truly dissociate deficits in attention and arousal, which are intertwined concepts, if overall arousal was compromising 5CSRTT error rate, we would additionally expect to see a decrease in the trial number attempted. Additionally, overall decreased arousal would lead to decreased breakpoint in our separate testing of motivation with progressive ratio. As arousal state should affect multiple measures of cognitive performance, and we observe deficits specific to error rate, this rather supports an attentional deficit over general arousal deficit. Together, these data point to adolescence as an important sensitive window for top-down ACA_VIS_ projection neurons to drive activity-dependent maturation of visual attentional behavior in adulthood.

### Adolescent top-down activity promotes local input integrity

We next set out to gain mechanistic insights into the later adult visual attentional deficits caused by transient adolescent suppression of top-down ACA_VIS_ neuron activity. To this end, we performed optogenetic interrogation of local inputs onto ACA_VIS_ neuron by patch-clamp recordings to examine the impact of adolescent chemogenetic suppression of ACA_VIS_ neuron activity to local excitatory connectivity which showed greater level during adolescence compared to adulthood in contrast to long-range excitatory inputs which did not differ between adolescence and adulthood. To virally express iDREADD-mCherry or mCherry in maturing ACA_VIS_ projection neurons, we infused retrograde Cre encoding virus in VIS and Cre-dependent iDREADD-mCherry or mCherry encoding virus in ACA around p12. We then virally introduced Flpo and FlpON/CreOff-ChR2 vectors into the ACA to express ChR2 selectively in Flpo-expressing ACA neurons yet actively avoid labeling ACA_VIS_ neurons expressing Cre at p52-p80. Adolescence iDREADD and adolescence mCherry groups experienced CNO injections during adolescence (p25-34) where adolescence mCherry group was used as a control both for the effect of CNO and surgery/injections. Late-adolescent iDREADD group experienced CNO from p45 to p54 to test the temporal specificity of ACA suppression **(Figure 6a)**. In all experimental groups, patch-clamp recordings were performed from ACA_VIS_ neurons in acute ACA slices obtained from adult mice (p85.7+/-2.3 in average (p72-107)). The spread and intensity of YFP signal of analyzed slices show comparable levels of ChR2 expression between groups (**Figure 6b, c:** see details in methods section).

**Figure 6:**
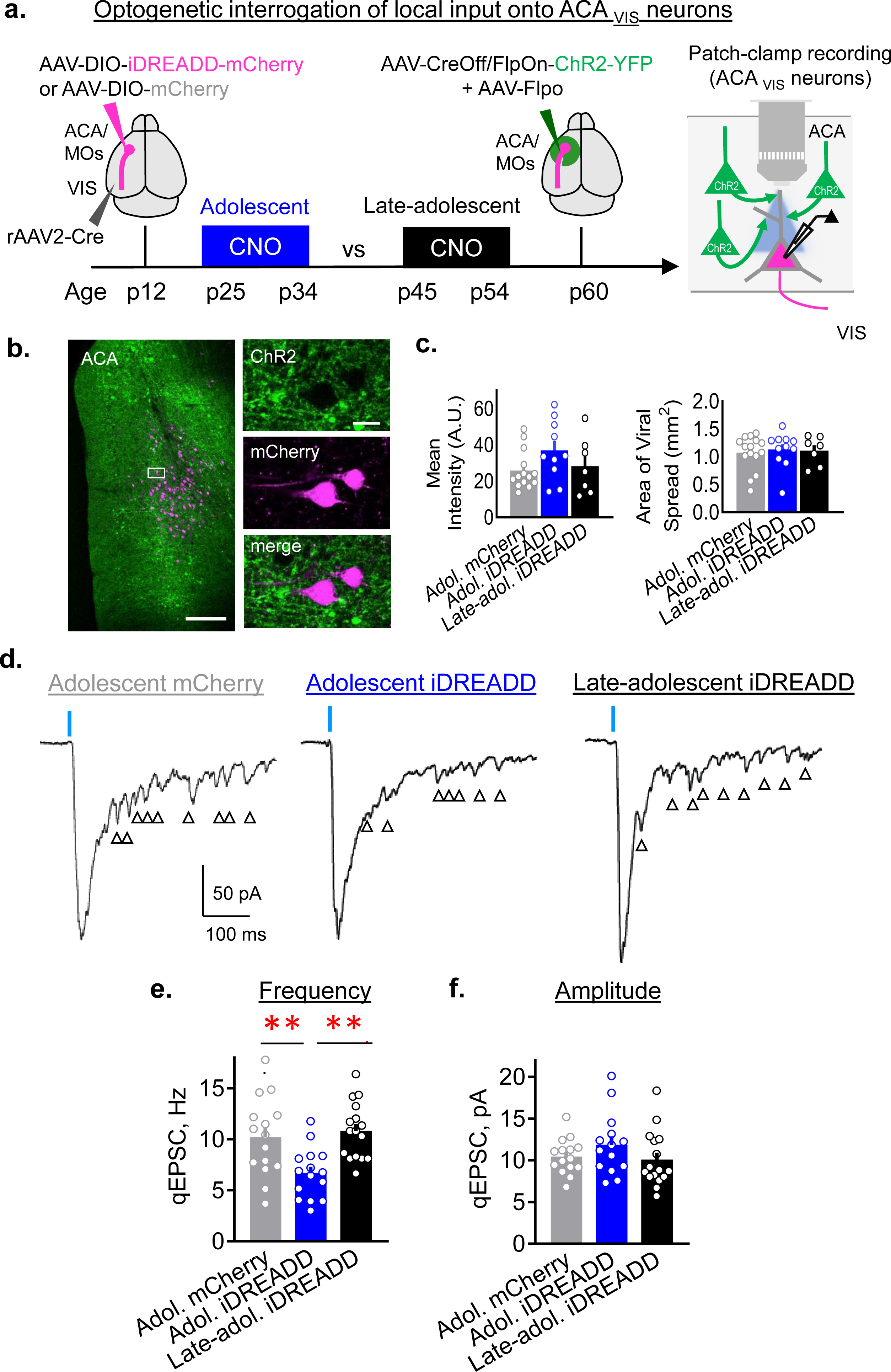
Suppression of adolescent ACA_VIS_ neuron activity results in their excessive loss of local excitatory inputs. **a**. Experimental approach to test the impact of ACA_VIS_ neuron activity during adolescent or late-adolescent period on maintaining local excitatory input drive onto ACA_VIS_ neurons into adulthood through optogenetic interrogation of local excitatory input onto ACA_VIS_ neurons. (left) RetroAAV2-cre was injected into the VIS, while AAV-DIO-hM4D(Gi)-(iDREADD)-mCherry or AAV-DIO-mCherry was injected into ACA at p12 to selectively express iDREADD-mCherry or mCherry in ACA_VIS_ neurons by p25. (middle) Mice were injected with 10 mg/kg CNO in normal saline twice a day from p25 to p34 in adolescent iDREADD and adolescent mCherry groups, or from p45 to p54 in late-adolescence iDREADD group. Additional injections of AAV-Flpo and AAV-CreOff/FlpOn-ChR2-YFP in ACA at p60.4±2.2 in average (p52-p80) enable ChR2-GFP expression in all but ACA_VIS_ neurons in ACA. (right) Slice patch clamp recordings were performed from ACA_VIS_ neuron at p85.7±2.3 in average (p72-107) while optogenetically activating ChR2-expressing local excitatory inputs. **b**. (left) Representative image showing ChR2-YFP expressing local inputs (green) and mCherry expressing ACA_VIS_ neurons (magenta) in ACA. (right) High magnification of boxed area showing mCherry+ACA_VIS_ neuron does not express ChR2-YFP and is surrounded by ChR2+ terminals. Scale bars: 200 μm on the left and 20 μm on the right. Experimental images were obtained from 24 mice with similar results obtained. **c**. The quantification of the spread and intensity of the YFP signal showed no significant differences among groups (one-way ANOVA, area: *F*_2,30_ = 1.334, *P*=0.8962, mean intensity: *F*_2,28_ = 1.334, *P*=0.1612, n=13 slices from 9 biologically independent mice for adolescent iDREADD. n=15 slices from 10 biologically independent mice for adolescent mCherry, and n=7 slices from 5 biologically independent mice for late-adolescent iDREADD). **d**. Examples of light-evoked EPSC in ACA_VIS_ neurons in the presence of strontium to produced delayed asynchronous release of neurotransmitters for each groups. Arrows indicate detected qEPSCs. **e-f**. Averaged qEPSC **(e)** amplitude and **(f)** frequency for adolescent iDREADD, adolescent mCherry and late-adolescent iDREADD groups (frequency: one-way ANOVA *F*_2,43_ =8.010, \*\**P*=0.0011, post hoc Tukey test \*\**P*_Adol.iDREADD-Adol.mCherry_=0.009, \*\**P*_Adol.iDREADD-Lateadol.iDREADD_=0.002; mean amplitude: one-way ANOVA, *F*_2,43_ =1.352, *P*=0.2697, n=15 cells from 4 biologically independent mice for adolescent iDREADD. n=15 cells from 4 biologically independent mice for adolescent mCherry, and n=16 cells from 4 biologically independent mice for late-adolescent iDREADD). Data in c, e-f are presented as mean +/- s.e.m. See related Supplementary Figures 8-9. Source data are provided as a Source Data file.

To assess consequence of developmental suppression of ACA_VIS_ neuron activity on local excitatory drive onto ACA_VIS_ neurons in adulthood, we recorded quantal EPSCs in the presence of strontium (Sr^2+^, 3 mM), TTX, 4-AP and picrotoxin in a bath (**Figure 6d**). Optogenetic stimulation of local inputs onto ACA_VIS_ projection neurons revealed decreased frequency of quantal synaptic events in adolescent iDREADD animals in comparison to adult iDREADD or adolescent mCherry groups following the light stimulation (**Figure 6d**). No differences were found in the amplitude of quantal events (**Figure 6d**). We also did not observe difference in paired-pulse ratio **(Supplementary Figure 8)**. These results strongly suggest that adult visual attention deficit caused by adolescent suppression of top-down ACA_VIS_ neuron activity parallels the excessive reduction of local excitatory inputs onto ACA_VIS_ neurons.

Next, to examine a structural correlate of our findings, we virally introduced a bright monomeric fluorescent protein (mHyperGFP1) that allows visualization of dendritic spines without immunohistochemical enhancement (Campbell et al unpublished) together with iDREADD selectively in ACA_VIS_ neurons **(Supplementary Figure 9a)** and suppressed activity in iDREADD expressing neurons by CNO administration during adolescence from p25 to p34 **(Supplementary Figure 9b)**. Injections volumes for DREADD and mHyperGFP1 were optimized so that some mHyperGF1P labelled neurons were co-infected with inhibitory DREADD (mCherry+), and thus allowed for within animal comparison with DREADD negative (mHyperGFP+ mCherry-) ACA_VIS_ neurons **(Supplementary Figure 9c and Supplementary 9d)** while also ruling out any potential DREADD independent CNO effect ^45^. Adolescent chemogenetic suppression produced an excessive decrease in adult total spine density mCherry+ suppressed ACA_VIS_ neurons compared to mCherry-control neurons selectively in proximal dendrites **(Supplementary Figure 9e)**. Of note, a previous study demonstrated that proximal dendrites of layer V/VI neocortical pyramidal neurons preferentially receive local inputs from surrounding layer II/III neurons ^46^, while different long-range inputs target select dendritic locations across dendrites ^46,47^. Observed loss of spines in proximal dendrites is consistent with the optogenetics-assisted functional data showing an excessive loss of local excitatory drive onto ACA_VIS_ projection neurons **(Figure 6)**. While future investigations are warranted to examine whether the impact of adolescent suppression of ACA_VIS_ neuron activity is limited to local excitatory inputs or also extends to long-range inputs, our study collectively suggests that adolescent top-down ACA_VIS_ projection neuron activity is required to maintain their local excitatory inputs and support adult visual attentional behavior.

## Discussion

In this study, we examined the maturation of frontal top-down corticocortical projection neurons, and demonstrated an adolescence as a time of elevated local excitatory input drive and heightened activity of top-down projection neurons prior to a transition into an adult state with increased long-range contribution through local circuit pruning. Transient perturbation of top-down projection neuron activity at this adolescent window produced long-lasting attentional behavior deficits in adulthood and is accompanied by excessive loss of local inputs. Collectively, our data suggests that heightened adolescent activity is essential for a frontal top-down projection to maintain local inputs and establish attention in adulthood **(Figure 7)**.

**Figure 7.**
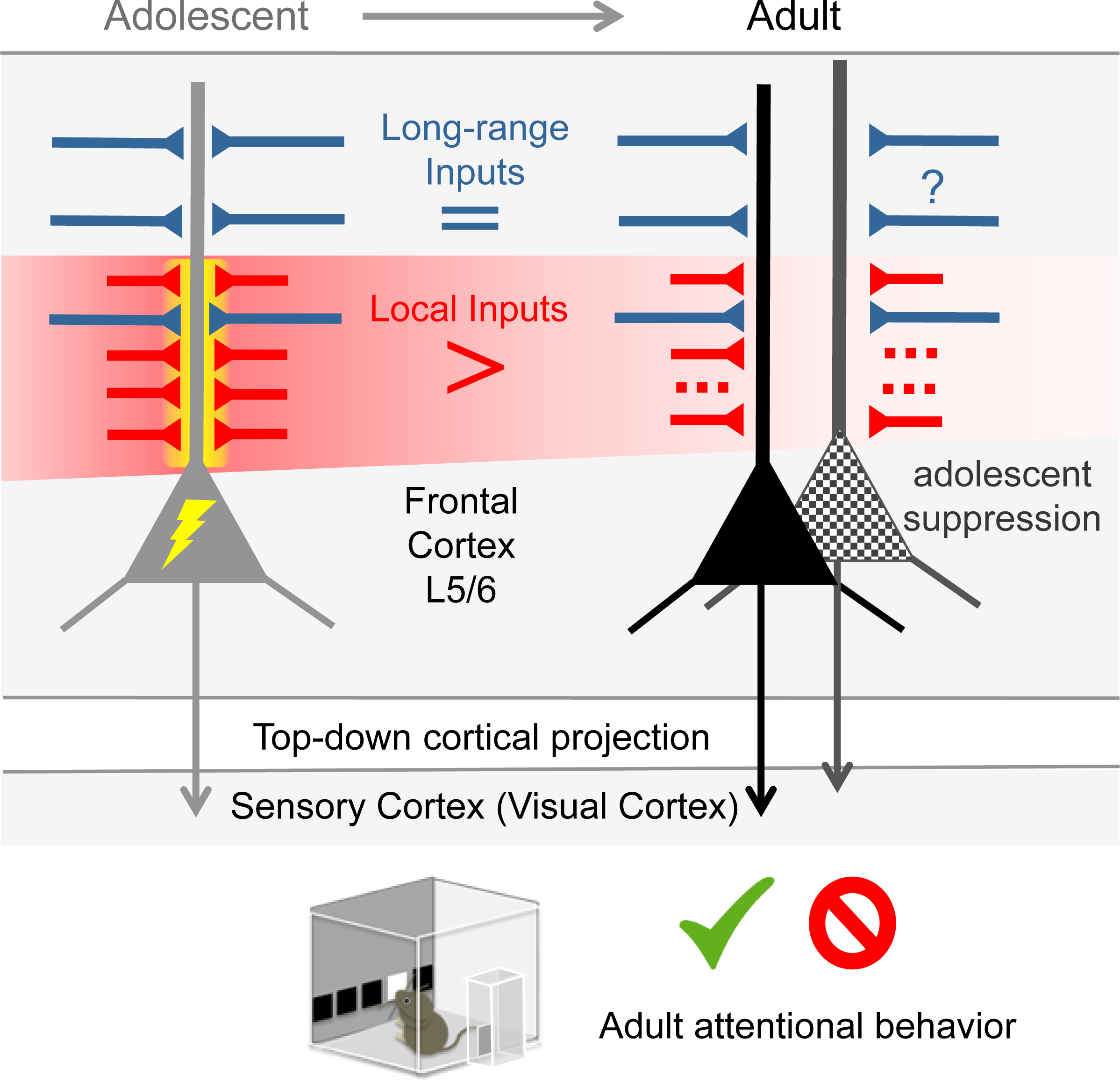
Summary: Adolescent period of top-down circuit maturation. Frontal top-down ACA_VIS_ projection neurons receive heightened local excitatory inputs (red) and show greater activity (yellow lightning bolt) during adolescence compared to adulthood, while long-range inputs (blue) are already established prior to adolescence. These developmental changes lead to a shift in the balance of local/long-range input on ACAvis neurons between adolescence and adulthood. Chemogenetic perturbation of top-down activity during this adolescent window of heightened excitatory and local connectivity (no lightning bolt) results in failed maintenance of local connectivity and attentional deficits in adults (top-down neuron underwent adolescent silencing is labeled with black dots).

Our circuit-specific patch recording, rabies input mapping, and optogenetic interrogations collectively identified a heightened local excitatory drive onto ACA_VIS_ projection neurons during adolescent period compared to adulthood. A postnatal waxing and waning of synaptic contacts is stereotypical across several cortical regions ^*48,49*^, but how this developmental event impacts input-output relationship was poorly characterized. This study relates such observations to circuit rewiring for a corticocortical projection and suggests that this process is driven by local events. These findings also provide a conceptual basis for considering similar phenomenon observed in human frontal cortex, in which synaptic density heightens during childhood ^*50,51*^. While the developmental changes revealed by this study were restricted to local excitatory input, we found a relative impact of local input pruning on long-range circuitry. Forming long-range synapses is more challenging and energetically intensive than establishing local connections in which processes can more easily extend and retract ^52-54^. The identified reduction in local drive may serve as a more efficient and effective manner of increasing the contribution of long-range connectivity. Further, as local inputs tend to preferentially target proximal dendrite ^46^, removal of irrelevant local circuits may aid somatic transmission from some of long-range inputs targeting distal dendrites. We propose a model that this post-adolescent shift in balance between local and long-range inputs is a key developmental milestone for top-down projection to effectively mediate attentional behavior. Future studies are warranted to examine to what extent these findings in ACA_VIS_ projection neurons are applicable to other prefrontal neurons located in different layers and/or projecting to different brain regions.

What is the purpose of heightening top-down neuron activity and associated greater local inputs if they are eventually diminished? At the circuit level, top-down neurons that experienced adolescent chemogenetic suppression showed over-reduction of local excitatory inputs **(Figure 6)** and total spine density on proximal dendrites in adulthood **(Supplementary Figure 9)**. We therefore speculate that adolescent heightened activity of top-down projection is required to allow robust activity-dependent post-adolescent synaptic refinement from a pool of enough number of local inputs to maintain adequate local connectivity into adulthood. Based on previous reports examining the role of activity in circuit formation, we speculate that highly-active local connectivity is stabilized, while insufficiently active contacts are eliminated or fail to be maintained ^55 56^. Our findings also suggest that physiological developmental reduction local connectivity onto ACA_VIS_ neurons following adolescence into adulthood is likely mediated by distinct mechanisms and requires future investigations. It is also important to examine in future studies if adolescent ACA_VIS_ projection neuron activity can additionally impact downstream ACA_VIS_ projection outputs onto VIS neurons. At the behavioral level, our study demonstrates that disruption of frontal top-down projection neuron activity during adolescence affects adult attentional behavior **(Figure 5)**. We speculate that an excessive local pruning, as produced by adolescent chemogenetic suppression may fail to effectively drive adult ACA_VIS_ neurons when attentional adjustment is necessary. Heightened activity of ACA_VIS_ neurons in conjunction with an elevated excitatory local input drive at the adolescent period may be therefore essential for the maturation of top-down neurons and associated connections within the frontal cortex to play a role in attentional processing and sensory cue-guided actions ^20,57,58^ as well as to effectively control distal sensory cortical neurons^9,10^. Future studies using additional cognitive tasks is necessary to examine the generalizability and selectivity of adolescent ACA_VIS_ activity manipulations to cognitive behaviors.

Our study provides an experimental support to a hypothesized framework that disrupted local vs long-range balance in frontal cortex may underlie cognitive deficits in psychiatric disorders ^59-61^, and stresses a need for future studies to examine the balance between local and long-range connectivity across development in the context of psychiatric disorders. Our transient adolescent manipulation produced deficits later in life, also points to a possible developmental mechanism for deficits in corticocortical functional connectivity, and attentional function. Disruption of circuit activity during the essential adolescent period could decrease frontal dendritic spine density **(Supplementary Figure 9)**, as is observed in post-mortem studies of schizophrenia and other neuropsychiatric disorders ^62,63^. Our study also proposes that failure to achieve this developmental milestone of activity-dependent local input refinement may produce attention deficits. Heterogeneous high-risk gene variations or environmental insults at this critical developmental stage could converge on failures of proper circuit rewiring and thus produce common functional deficits—as is observed across attention deficits in autism, schizophrenia, and ADHD ^2,14-16^. Clinical approaches that protect and elicit sufficient excitatory drive and top-down projection activity during this adolescent sensitive period could ameliorate visual attention deficits in neurodevelopmental disorders. Further, interventions promoting activity-dependent spine formation may have therapeutic value for adult deficits. Interrogating these possibilities in mouse models carrying these circuit deficits would provide valuable preclinical insight.

## Methods

### Animals

Male C57Bl/6 mice (Charles River) were group-housed in standard laboratory cages in a temperature (72 degrees Fahrenheit)- and humidity (30-70%)-controlled vivarium with a 12:12 light/dark cycle. Food and water were provided *ad libitum* throughout the experiment. Only male mice were used for experiments. To breed experimental mice, both sex were used. Adolescent mice were shipped at 10 days after birth (P10) and weaned at P21. Adult mice were shipped at P56. Experiments were performed when mice were between P12 and 6 months old. During behavior testing, mice were allowed access to water for 2 hr each day and maintained approximately 85-90% of respective baseline free feeding weight. All animal protocols were approved by IACUC at the Icahn School of Medicine at Mount Sinai.

### Drugs

Clozipine-N-Oxide dihydrochloride (CAS2250025-93-3, Tocris Biosciences) was injected intraperitoneally (i.p.) two times a day at 10mg/kg in sterile saline for 10 days from p25 to p34, or from p45 to p54.

### Stereotactic Surgery

Mice were anesthetized with 2% isoflurane and head-fixed on in a mouse stereotactic apparatus (Narishige International USA Inc.) equipped with a heating pad. Mice were isolated post-surgery until fully-awake and immediately returned to their home cage. VIS injection sites relative to lambda are: AP +0.0 mm, ML +3.0 mm, DV −0.4 mm; AP +0.1 mm, ML +2.85 mm, DV −0.4 mm; AP +0.1 mm, ML +3.15 mm, DV −0.4 mm. ACA injections sites relative to bregma are: AP +1.7 mm, ML +0.2 mm, DV −0.7 mm; AP +1.1 mm, ML +0.2 mm, DV −0.7 mm; AP +0.4 mm, ML +0.2 mm, DV −0.7 mm. MOs injection sites relative to lambda are: AP +0.5 mm, ML +0.4 mm, DV −1.0 mm; AP +0.0 mm, ML +0.4 mm, DV −1.0 mm; AP −0.5 mm, ML +0.4 mm, DV −1.0 mm. For patch-clamp recordings, 500 nl of fluorescent retrobeads (Lumafluor, Inc) were injected into the VIS 4 to 6 days prior to patch recordings to allow visualization of top-down projection neurons. 350 nl of AAV5-hSyn-eGFP-WPRE (University of Pennsylvania Viral Core) was infused into VIS for the basic characterization of ACA_VIS_ projection neurons. For rabies-mediated monosynaptic input mapping, 500 nl of a 1:1 volume mixture of AAV8-CA-FLEX-RG and AAV8-EF1a-FLEX-TVAmCherry (University of North Carolina Viral Core) were injected unilaterally in the ACA and 500 nl of CAV2-Cre (Institut de Genetique Moleculaire de Montpellier) were injected in VIS. For somatic introduction of rabies, 2.5 weeks later, 500 nl of pseudotyped EnvA RVdG-GFP (Salk Institute Viral Core) was injected into the ACA for local axonal uptake of rabies or VIS for distal axonal uptake of rabies. Mice were euthanized 7 days later. For behavioral studies, 500 nL of AAV8-DIO-hM4D(Gi)-mCherry and 500 nl CAV2-Cre were injected in the ACA/MOs and VIS, respectively. For the dendritic spine study, a total of 500 nl of AAV8-DIO-hM4D(Gi)-mCherry (University of North Carolina Viral Core) and AAV1-EF1a-DIO-hyperGFP2-WPRE (Custom Prep, Boston Children’s Hospital) at a ratio of 2.5:1 was injected in the ACA and 500 nl CAV2-Cre (Institut de Génétique Moleculaire de Montpellier) in VIS sites at p12. For optogenetics-based patch clamp recordings, rAAV2-Ef1a-mCherry-IRES-Cre (Addgene) was injected into VIS while AAVDJD/hSyn-CreOff/FlpOn-ChR2-YFP and AAV5/EF1a-FLPO-WPRE (both from University of North Carolina Viral Core) were injected in ACA at p11-15 for adolescent group and at p38-58 for late-adolescent group. For the developmental experiments with iDREADD, rAAV2-CAG-Cre-WPRE (Boston Children’s Hospital) was injected into VIS while AAV8-hSyn-DIO-hM4D(Gi)(iDREADD)-mCherry or AAV8-hSyn-DIO-mCherry (Addgene) was injected into ACA at p12-13. Additional injections of AAV5/EF1a-Flpo and AAVDJD-hSyn-CreOff/FlpOn-ChR2-YFP in ACA were at p52-p80. For the *in vivo* electrophysiology experiment where ACAvis neurons were optogenetically tagged and recorded, a virus expressing channelrhodopsin-2 (rAAV2-Syn-ChR2(H134R)-GFP: Addgene) was injected into the VIS.

### Tissue preparation and Histology

Animals underwent transcardial perfusion with phosphate-buffered saline (PBS) followed by 4% paraformaldehyde (PFA) in PBS. Brains from rabies input mapping and DREADD validation were dissected, postfixed in 4% PFA for 3 h, and cryoprotected in 30% sucrose in PBS for 24-48 h before embedding in Optimum Cutting Temperature (OCT, Tissue Tek) and sectioning into 35 µm-thick coronal slices using a cryostat (CM3050, Leica). For rabies input mapping, free-floating sections were washed in PBS before undergoing immunohistochemistry by 1hour incubation in blocking solution (1% bovine serum albumin in 0.1% Triton X-100 in PBS) followed by overnight incubation with rabbit anti-eGFP antibody (Life Technologies # A-11122; 1;1000) at room temperature. Sections were washed 3 times with blocking solution followed by incubation with Alexa 488-conjugated goat anti-rabbit IgG (H+L) (Life Technologies:# A-11034; 1:5000) for 2.5 h, washed 3 times in blocking solution, and mounted onto slides with DAPI Fluoromount-G (Southern Biotech). For spine analysis, brains were post-fixed in 4% PFA overnight and coronally sectioned at 75 µm on a Vibratome (Leica VT1000 S). All slices were then mounted on Superfrost Plus slides using Fluoromount-G mounting medium (Southern Biotech) and coverslipped. DREADD injections were validated using a LSM780 confocal microscope (Zeiss) in reference to the Paxinos and Franklin mouse atlas. For the viral spread validation of mice used for behavioral experiments, mice completed behavior testing underwent transcardial perfusion with PBS followed by 4% PFA in PBS, with overnight postfixation. Brains were coronally sectioned on a Leica CM3050 S cryostat (Leica Microsystems, Buffalo Grove, IL) at 60 μm for immunohistochemistry. Slices were collected at nine specific Bregma areas (2.10, 1.70, 1.18, 0.74, 0.14, −0.34, −0.82, −1.34, −1.82) to analyze the anterior-posterior spread of virally infected cells. Images were acquired using the EVOS FL Imaging System (Thermo Fisher Scientific). Slices were imaged using a 4x lens for mCherry and to ensure consistency across imaging sessions, power was set at 100%. Using Microsoft Excel v. 14.7.3 (Microsoft), all groups were assigned five levels representing increasing levels of signal intensity. The first level (L1) represented the minimum number of mice showing signal in a given area, namely n=1. The second level (L2) represented the 25th percentile of mice with overlapping signal in a given area, the third level (L3) represented the 50th percentile, and the fourth level (L4) represented the 75th percentile. The fifth level (L5) represented the maximum number of mice showing overlapping expression in a given area, namely the total number of mice in the group. Using the GNU Image Manipulation Program v. 2.10.8 (GIMP), areas with mCherry signal were delineated on templates taken from the Paxinos and Franklin mouse atlas. Signal intensity was determined by the quartile and the number of mice with overlapping viral expression in a given area.

### Slice Electrophysiology

Animals were decapitated under isofluorane anesthesia. Brains were quickly removed and transferred into ice-cold artificial cerebrospinal fluid (ACSF) of the following composition (in mM): 210.3 sucrose, 11 glucose, 2.5 KCl, 1 NaH_2_PO_4_, 26.2 NaHCO_3_, 0.5 CaCl_2_, and 4 MgCl_2_. Acute coronal slices of ACA (300 μm) contained both hemispheres. Slices were allowed to recover for 40 min at room temperature in the same solution, but with reduced sucrose (105.2 mM) and addition of NaCl (59.5 mM). Following recovery, slices were maintained at room temperature in standard ACSF composed of the following (in mM): 119 NaCl, 2.5 KCl, 1 NaH_2_PO_4_, 26.2 NaHCO_3_, 11 glucose, 2 CaCl_2_, and 2 MgCl_2_.

Slices were visualized under upright differential interference contrast microscope equipped with high power LED-coupled light source used for the identification of fluorescently-labeled cells and optogenetic stimulation (Prizmatix). Patch clamp recordings were made from ACA_VIS_ projection neurons fluorescently labeled with retrobeads or mCherry. For voltage clamp recordings, borosilicate glass electrodes (5–7 MΩ) were filled with the internal solution containing (in mM): 120 Cs-methanesulfonate, 10 HEPES, 0.5 EGTA, 8 NaCl, 4 Mg-ATP, 1 QX-314, 10 Na-phosphocreatine, and 0.4 Na-GTP (pH 7.25, 295 mOsm). Current clamp recordings were obtained with the internal solution containing (in mM): 127.5 K-methanesulfonate, 10 HEPES, 5 KCl, 5 Na-phosphocreatine, 2 MgCl_2_, 2 Mg-ATP, 0.6 EGTA, and 0.3 Na-GTP (pH 7.25, 295 mOsm). Data were low-pass filtered at 3 kHz and acquired at 20 kHz using Multiclamp 700B (Axon Instruments) and pClamp 10 v. 10.6.2.2 (Molecular Devices). Neurons were included in the analysis if input resistance, series resistance and membrane potential did not change more than 10% during the course of recordings. Miniature inhibitory and excitatory postsynaptic currents were isolated by holding the neuron at the reversal potential for excitatory or inhibitory currents. mIPSC and mEPSC were recorded in the presence of TTX (1 μM; Abcam) in the bath solution to block action potentials. After acquisition of a stable baseline (10 min), miniature postsynaptic events were recorded 3-5 min for each potential.

For iDREADD validation spontaneous firing rate was measured in a current clamp mode. Recordings were performed in normal ACSF with a depolarizing current of 1.6±0.04 times rheobase injected to the cell. In order to establish steady-state firing activity of recorded neurons, we slowly increased the depolarizing current over the course of 2-3 minutes to elicit low level of sustained spontaneous firing. The recordings were initiated after the adjustment of the depolarizing current. Firing rate was quantified as the average instantaneous firing frequency measured over 3 min before and following CNO (10 μM) perfusion.

To selectively activate local inputs to frontal ACA_VIS_ projection neurons, we used slices being introduced with ChR2-GFP construct in ACA using the intersectional viral approach ^31^. ChR2 was activated with a 0.25-0.5 ms light pulse delivered through the objective and synchronized with electrophysiological recordings. At the beginning of each experiment an input/output curve was established for each neuron to determine the intensity of the stimulation that evoked EPSC reliably at the top of the curve to ensure activation of all ChR2-expressing terminals within the field of view. The power of light stimuli was quantified using an optical power meter (Thorlabs). The range of intensities at the tissue level was 0 to 3.9 mW/mm^2^. For all optogenetic experiments we set the intensity of the stimulus at the plateau level where additional increase in the light intensity didn’t affect the amplitude of the light-evoked responses. Short-term dynamics were tested with twin pulses separated by 100, 250 and 500 ms. Stimuli were given every 20 sec and four responses were averaged at each interpulse interval. Paired-pulse recordings were conducted in standard ACSF. To limit the impact of disynaptic excitation, paired pulse ratio was determined as a ratio of the slope of EPSC_2_ to the slope of EPSC_1_. To isolate monosynaptic connections, eEPSCs were recorded in the presence of TTX (1 μM), 4-AP (100 μM) and picrotoxin (100 μM). Quantal EPSC were measures in ACSF where CaCl_2_ was replaced with 3 mM SrCl_2_ in addition to TTX, 4-AP and picrotoxin. Quantal EPSCs were detected and analyzed using MiniAnalysis v. 6.0.7 (Synaptosoft Inc.). The detection threshold for events was set at 2x root mean square noise measured before stimulation and quantified by Noise Analysis tool built in MiniAnalysis. Evoked quantal EPSCs were detected in 30-400 msec from the stimulus onset, acquired for not less than 3 sweeps and averaged.

To assess consistency of ChR2-GFP expression across preparations, the spread and intensity of YFP expression was quantified for slices used in patch-clamp experiments. Following the recording, slices were fixed in phosphate buffer saline containing 4% PFA and stored at 4°C. Slices were subsequently washed three times for 5 min each in PBS and mounted on microscope slides (Brain Research Laboratories) using Fluoromount-G mounting medium (Southern Biotech). Images were obtained with an EVO FL Imaging System equipped with 4x objective to visualize entire ACA with the adjacent cortical areas simultaneously. Images were analyzed using ImageJ32 v. 1.51j8 software (NIH). Images were thresholded at mean intensity plus two standard deviation of the background fluorescent signal. The background fluorescent signal was calculated as an intensity of fluorescence in motor/somatosensory areas lacking YFP expression. The spread and intensity of YFP expression were quantified as the area and mean intensity of YFP signal respectively. Recordings were excluded from the analysis if obtained from slices in which the YFP signal deviated more than two standard deviation from the average intensity or area. This exclusion criteria led to exclude 1 slice from adolescent iDREADD group. This allowed for comparisons of recordings obtained from slices with similar levels of ChR2 expression in ACA.

### Monosynaptic Input Mapping

Samples were imaged using a LSM780 confocal microscope (Zeiss). All slices around the ACA injection regions were stained for starter cell analysis. One out of every 8 serial sections was stained for input analysis. ImageJ32 v. 1.51j8 software (NIH) were used to process and analyze all images. Starter cells were defined as both mCherry+ (>50% of soma pixels above 2 SD the mean image mCherry fluorescence intensity) and eGFP+ (>50% of soma pixels above 3SD the mean image GFP fluorescence intensity). Input cells were defined as eGFP+ (>50% of soma pixels above 3 SD the mean image eGFP fluorescence intensity). To determine the brain region location for all input and starter cells, all brain slices were registered to corresponding Allen Brain Institute coronal maps using anatomical landmarks visualized by DAPI counterstaining and tissue autofluorescence. In a small minority of cases, assignment of input neurons to specific brain nuclei may be approximate if eGFP+ cell bodies were located on borders between regions, or when anatomical markers were lacking between directly adjacent regions. However, quantitative analyses of input tracing results were performed on anatomical classifications that were at least one hierarchical level broader (as specified by the Allen Brain Atlas) than the discrete brain regions input cells were assigned to. Notably in many of these cases, directly adjacent nuclei fall together into the same hierarchical group. Both eGFP+ input neurons and eGFP+ mCherry+ starter cells were manually counted using the Cell Counter plug-in in ImageJ. For the input analysis, we normalized the total number of input neurons in each brain to the total number of starter cells because the efficiency of rabies uptake by helper cells differed slightly across animals. The investigator performing these analyses remained agnostic of the group identities of animals through the duration of this analysis.

### *In vivo* Extracellular Electrophysiology

Preparatory surgery leading to, and recording itself was performed initially under Nembutal/chlorprothixene anesthesia and then maintained with isoflurane^64^. Optogenetically tagged ACAvis neurons were identified using a laser (wavelength 473 nm, 1ms duration, 5 Hz) search stimulus emitted and delivered through the optic fiber (diameter 105 μm) coupled with a linear sixteen-channel silicon probes with 177 μm^2^ recording sites (NeuroNexus Technologies, Ann Arbor, MI) spaced 50 μm apart and oriented immediately above the ACA cortical surface. The power at the fiber-optic tip was approximately 10 mW. The signal detected from the probe was amplified and thresholded, and unit-sorted using an OmniPlex Neural Recording Data Acquisition System A (Plexon). A template online sorting method was used to capture spikes as units. A PCA cluster-based spike shape template-based sorting method was used. Visually identifiable subsets of waveforms form clusters on 2D PCA space. Spike shape templates were selected by manually drawing selection contours around PCA clusters. The selected spikes were then automatically averaged and used to create templates, which were then used to sort incoming baseline spikes of similar shape and placement in PCA space into units. After sorting for optogenetically responsive units (ACAvis neurons expressing ChR2), the optogenetic stimulus was switched off. Then baseline spike activity was recorded from the sorted units for 3 mins followed by 1min of recordings upon turning on a laser for optogenetic tagging. To ensure single-unit isolation, the waveforms of recorded units were further examined using Offline Sorter (Plexon). To analyze the electrophysiology data, firing rate was computed during 3 min laser off session mentioned above by a MATLAB R2018b (MathWorks) script established in a published study ^64^. Only sorted units whose spike width time (trough-to-peak time) is more than 400 μs were included in the analysis, and were further separated to optgenetically tagged ChR2 (+) ACAvis neurons if they showed time-locked spiking response within 2 ms upon laser pulse stimulation, and ChR2 (-) ACA neurons for the rest of units. For each animal, single units were recorded in each of the AP −0.3 mm, 0.0 mm, +0.3 mm, +0.6 mm, ML ±0.25 mm, DV −1.0 mm vertical penetrations across ACA.

### Five-Choice Serial Reaction Time Task (5CSRTT)

Behavior was conducted in black plastic trapezoid Bussey– Saksida touch-screen chambers (walls 20-cm high × 18-cm wide (at screen-magazine) × 24-cm wide (at screen) × 6-cm wide (at magazine) (Lafayette Instrument). Stimuli were displayed on a touch-sensitive screen (12.1-inch, screen resolution 600 × 800) divided into five response windows by a black plastic mask (4.0 × 4.0-cm, positioned centrally with windows spaced 1.0 cm apart, 1.5 cm above the floor) fitted in front of the touchscreen. Schedules were designed and data were collected and analyzed using ABET II Touch software v. 18.04.17 (Lafayette Instrument). The inputs and outputs of the multiple chambers were controlled by WhiskerServer software v. 4.7.7 (Lafayette Instrument). Before training on 5CSRTT, mice were initially trained to touch the screen. Mice were acclimated to the chambers for 3 days in 30-min sessions. During this habituation phase, the food magazine was illuminated and diluted sweetened condensed milk (Eagle Brand) was dispensed every 40 s. Mice were then trained to touch the response windows. If the mouse touched the stimulus, the milk reward was delivered in conjunction with a tone and magazine light. Touches to non-stimuli had no consequence. After reaching criterion on this phase (20 touches in 30 min for 2 consecutive days), mice moved onto 5CSRTT training phase. Mice were tested 5 days a week, 100 trials a day (or up to 30 min). Each trial began with the illumination of the magazine light. When the mouse made a nose poke in the food magazine, the stimulus was delivered after an intertrial interval (ITI) period of 5 s. If a mouse touched the screen during this ITI period, the response was recorded as premature and the mouse was punished with a 5-s time-out (house light on). After the time-out period, the magazine light illumination and house light switch off signaled onset of the next trial. After the ITI period, a stimulus appeared randomly in one of the five response windows for a set stimulus duration (this varied from 32 to 2 s). A limited-hold period followed by the stimulus duration was 5 s, during which the stimulus was absent but the mouse was still able to respond to the location. Responses during stimulus presence and limited holding period could be recorded either as correct (touching the stimulus window) or incorrect (touching any other windows). A correct response was rewarded with a tone, and milk delivery, indicated by the illumination of the magazine light. A failure to respond to any window over the stimulus and limited-hold period was counted as an omission. Incorrect responses and omissions were punished with a 5-time-out. In addition, animals could make perseverative responses that are screen touches after a correct or incorrect response. Animals started at stimulus duration of 32 seconds. With a goal to baseline mice at a stimulus duration of 2 s, the stimulus duration was sequentially reduced from 32, 16, 8, 4, to 2 s. Animals had to reach a criterion (>50 trials, >80% accuracy, <20% omissions) over 2 consecutive days to pass from one stage to the next. After reaching baseline criterion with the 2 s stimulus duration (5 consecutive days), mice were challenged with an increased attentional demand by reducing the stimulus duration to 2, 1.5, 1, and 0.8 s (reduced stimulus test). They then underwent 4 days of testing. Attention and response control were assessed by measuring the following performance: percentage error (100 x (incorrect responses + omissions))/(correct responses + incorrect responses + omissions), percentage premature response (100 × premature responses/(omissions + correct responses + incorrect responses), percentage perseverative response (100 × perseverative responses /(correct responses + incorrect responses), latency to correct response, and latency to reward collection after correct choices.

### Progressive Ratio Task

Behavior was conducted following completion of the 5CSRTT in touch-screen chambers. In initial operant training, mice were trained to nose-touch the visual stimulus (white square, 100% luminance) at the center screen one time in each trial to receive a diluted sweetened condensed milk (Eagle Brand) reward as dispensed in the 5CSRTT. Touches to non-stimuli had no consequence. After reaching criterion of at least 30 completed trials in 30 mins, mice moved onto fixed ratio training. In fixed ratio training, mice were trained to nose-touch the visual stimulus two, three, then five times to get a reward. Mice were required to reach criterion for one day for fixed ratio of 2:1 and 3:1, and for three days for fixed ratio 5:1 before proceeding to progressive ratio training and testing. Task parameters were identical for progressive ratio training, except that completion of each trial, the number of touches required to receive a reward was incremented on a linear +4 basis (i.e. 1, 5, 9, etc.). If no stimulus response or magazine entry in the presence of a delivered reward was detected for 5 minutes, the session ended and the animal was removed from the chamber. Otherwise the session ended after 30 min. Mice had to achieve a stabile breakpoint defined as the number of target location responses in the last successfully completed trial for 2 days before proceeding to progressive ratio testing. In progressive ratio testing, mice performed the progressive ratio task using the same parameters as progressive ratio training, but were allowed 60 min to complete the session. Mice were baselined on Monday with progressive ratio training to ensure consistent performance and tested for four consecutive days. Breakpoint across the four days was averaged to give each mouse on breakpoint score.

### Dendritic Spine Imaging and Characterization

Images were acquired using an Upright LSM 780 Confocal microscope (Carl Zeiss). A non-biased, whole neuron approach was taken to characterize dendritic spines. Specifically, two neurons per animal were imaged on both the apical and basal dendrites. To qualify for spine analysis, whole neurons were selected that met the following requirements: (1) the neuron had to be filled and display spines at least up to 200 µm away from the soma on both apical and basal dendrites and (2) display no overlapping of dendritic processes with other neuron processes. For the chemogenetic study, mCherry+ and mCherry-neurons were defined as having >50% of soma pixels above 3 SD and below 2 SD the mean image mCherry fluorescence intensity, respectively. Images were taken of sections from two basal dendrites and two apical dendrites from each neuron and binned into 4 distance groups from the soma: 0-50 µm, 50-100 µm, 100-150 µm, and 150-200 µm. Dendritic segments were imaged using a 100X lens (numerical aperture 1.4; Carl Zeiss) and a zoom of 3.0. Images were taken with a resolution of 1024 x 300, pixel dwell time was 1.58 µm/s, and the line average was set to 8. Pixel size was 0.03um in the x–y plane and 0.01 µm in the z plane. To assure consistency of imaging across difference confocal sessions, power was consistently set to 3.0%, and gain was adjusted within the range of 600-800 units to achieve consistent light intensity values within the same set of neurons.

Images were deconvolved using a 3D resolution enhancement with AutoDeblur X3 (Media Cybernetics) and then run through the dynamic range filter in Neuron-Studio v. 2.26.16 ^65^. To detect dendritic spines from background noise accurately, puncta were only counted as spines if they moved into and out of the z-plane with the dendritic branch. Spine density was calculated as the total spine count/dendritic length. All imaging and analysis were all performed by an author who remained agnostic to the group conditions until the analysis was completed.

### Statistical Methods

Linear mixed modeling approaches using the packages LmerTest (v. 2.0.32), lme4 (v. 1.1.12), and lsmeans (v. 2.25) in the R programing language (v. 3.2.2) were user for analysis of starter cell distribution. Groups were modeled as fixed effects and animals were modeled as random effects (this factor considered independent and randomly sampled from their respective populations) to determine the impact of group on location of starter cell.

All other statistical analyses were performed using Prism v8 (Graphpad). In electrophysiological recordings differences in amplitude and frequency of postsynaptic currents as well as the spread and intensity of YFP expression were tested by a two-tailed unpaired *t*-test, or a one-way repeated measures analysis of variance (ANOVA) followed by post hoc Tukey’s multiple comparisons test, while the effect of CNO on membrane potential was determined with two-tailed paired t-tests. Paired pulse ratio analysis was tested by two-way repeated measures ANOVA. For input mapping and ratio comparisons, input numbers were first normalized to the number of starter cells within each animal before analyses. Input mapping analyses were completed by a two-way repeated ANOVA with brain regions as within-subjects factors, followed by post hoc Sidak’s multiple comparisons test. Input ratios and locally restricted rabies experiment were both analyzed using a two-tailed unpaired t-test. 5CSRTT data (error) were analyzed using a two-way repeated measures analysis of variance (ANOVA) with and stimulus duration (2, 1.5, 1, 0.8 second) as within-subjects factors, followed by Newman-Keuls multiple comparisons test. In the analysis of other 5CSRTT data and progressive ratio testing, an unpaired t-test was used. Dendritic spine data were analyzed with ANOVA tests with two-way repeated measures with distance from soma (0-50 µm, 50-100 µm, 100-150 µm, 150-200 µm) as within-subjects factors, followed by post hoc Sidak’s multiple comparisons test. All data are expressed as means ± SEM.

## Data availability

Source data are provided with this paper.

## Code availability

All scripts used to analyze or display the data are available upon reasonable request.

## Acknowledgements

This work was supported by National Institute of Health F30 MH111143 to E.N., R21 MH106919, R21 NS105119, R01 MH119523 to H.M., T32 DA007135 to M.P.D., Seaver Foundation to E.N. We thank Milo Smith for assisting statistical analysis, Annalee Tacuri for assisting histological analysis, Dr. Mark Baxter for his expertise on touch-screen behavior system, Dr. Yasmin Hurd and Dr. Michael Michaelides for their expertise on DREADD.

## Author Contributions

E.N., Y.G., H.K., and H.M. designed research; E.N., Y.G., H.K., M.S., A.L., K.N., G.T., S.I., K.C., S.L, M.P.D, J.B. performed experiments; E.N., Y.G., H.K., A.L., G.T., S.I., K.C., and H.M. analyzed data; P.R.H., R.L.C., H.M. supervised the research, E.N., Y.G. and H.M. wrote the manuscript with contributions from all other co-authors. The authors declare no conflicts of interest.

## Competing Interests

The authors declare no competing interests.

## Notes

### Competing Interest Statement

The authors have declared no competing interest.

