## Supplemental Figures for "Adolescent frontal top-down neurons receive heightened local drive to establish adult attentional behavior in mice"

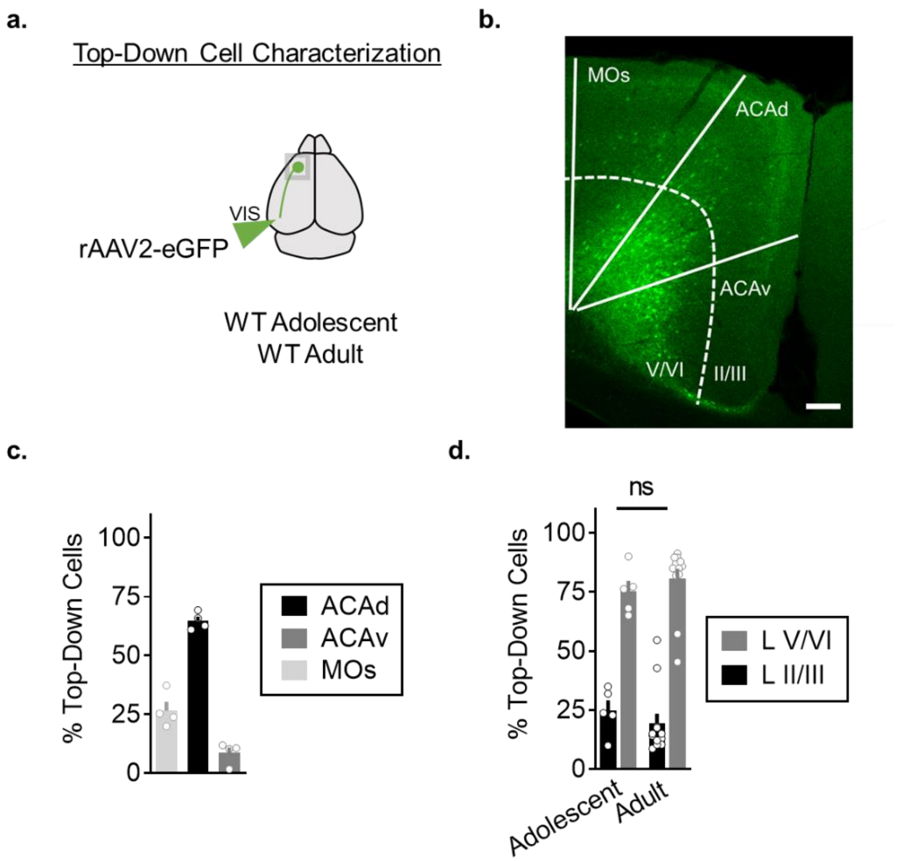


**Supplementary Figure 1. Characterization of frontal to visual cortical top-down projection neurons (related to Figure 1): a.** Schematic overview of retrograde viral labeling of projection neurons to primary visual cortex. Retroviral eGFP was injected into visual cortex and allowed to express for 7 days. **b.** Histologic sections were stained with Neurotrace Nissl staining to delineate cortical layers and total number of eGFP+ cell bodies were subsequently counted. Representative image of frontal cortical projection labeling and distribution among cortical layers (Scale bar =100 µm). Experimental images were obtained from 4 biologically independent mice, three images per mouse, with similar results obtained. **c.** Quantification of location percentage of projection neurons identified in frontal cortical regions revealed approximately 65% arising from dorsal anterior cingulate cortex (ACAd), 9% arising from ventral anterior cingulate cortex (ACAv), and 27% arising from secondary motor area (MOs) (n=4 biologically independent mice, 6 sections per mouse). **d.** Quantification of cortical layer distribution of frontal projection neurons to visual cortex in adolescent and adult mice showed approximately 75-81% arising from layers V/VI and 19-25% arising from layers II/III) (n=5 hemispheres from 3 biologically independent mice for adolescent, 12 hemispheres from 6 biologically independent mice for adult, 3 sections per hemisphere, two-tailed unpaired *t*-test, $t_{15}$=0.764, $P$=0.457). Data in c-d are presented as mean +/- s.e.m. Source data are provided as a Source Data file.

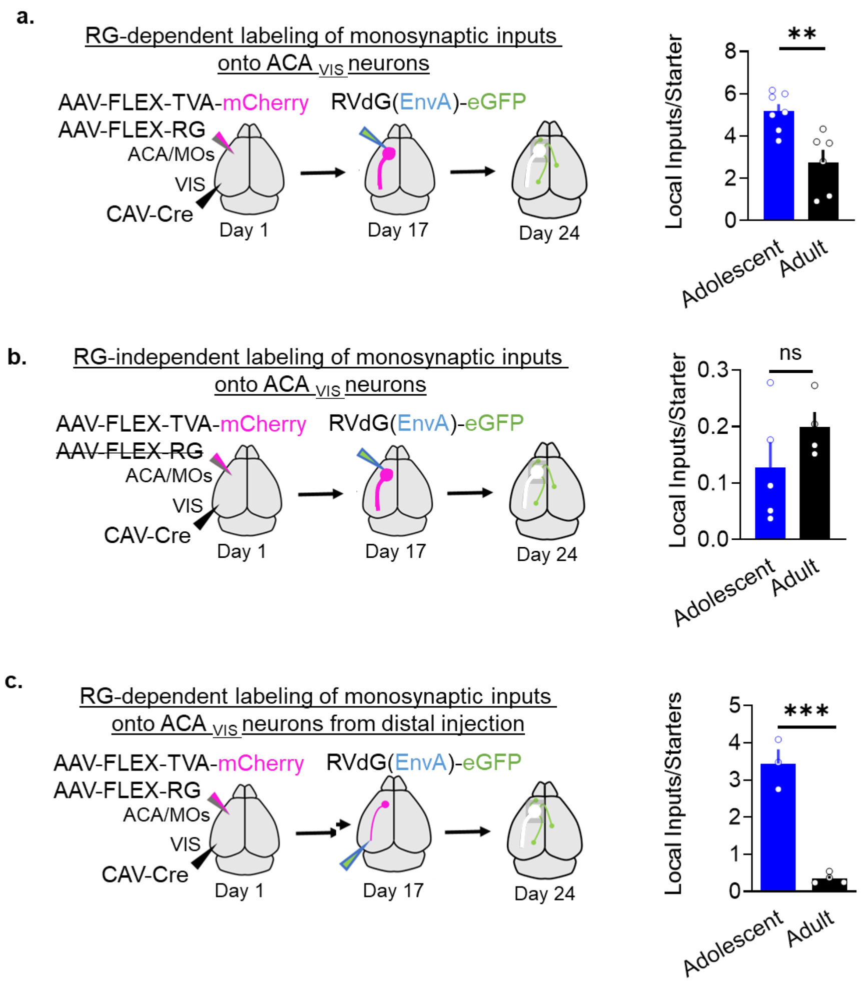


**Supplementary Figure 2. Control experiments for rabies virus-based mapping of inputs onto ACA$_{VIS}$ neurons (related to Figure 2): a**. When AAV-FLEX-TVA-mCherry was injected *together with* AAV-FLEX-RG followed by local RVdG(EnvA)-eGFP injection in ACA (as in Figure 2), the number on local inputs was higher in the adolescent group compared to the adult group (Two-tailed unpaired *t*-test, $t_{11}$=3.682, **$P$=0.0036, n=7,6 biologically independent mice). **b.** When AAV-FLEX-TVA-mCherry was injected *without* co-injection of AAV-FLEX-RG followed by local RVdG(EnvA)-eGFP injection in ACA, only a low amount of leak was detected in both groups (GFP+TVA- cells) were observed, which was not age-dependent (Two-tailed unpaired t-test, $t_7$=1.273, $P$=0.2436, n=5,4 biologically independent mice). **c.** When rabies virus was injected distally in the visual cortex to be taken up by ACAvis terminals to avoid local leak,  the number on local inputs was higher in the adolescent group compared to the adult group (Two-tailed unpaired *t*-test, $t_5$=9.224, ***$P$=0.0003, n=3,4 biologically independent mice). Data are presented as mean +/- s.e.m. Source data are provided as a Source Data file.

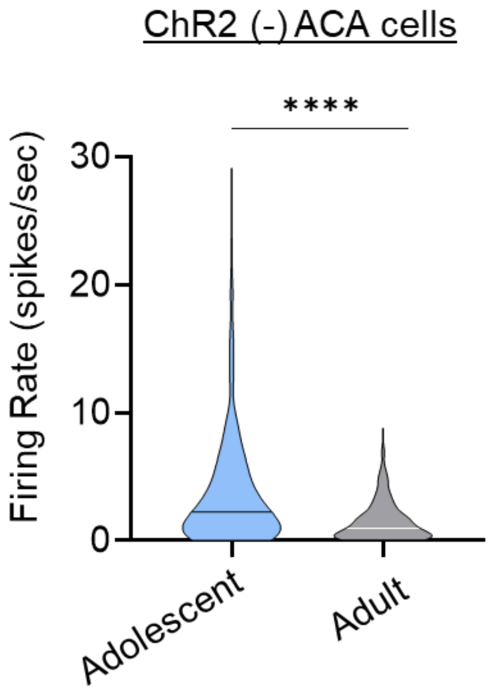


**Supplementary Figure 3. Firing activity of ChR2 (-) ACA neurons *in vivo* (related to Figure 4):** *in vivo* electrophysiological recordings of non-optically tagged ACA neurons in adolescent and adult mice in which retrograde AAV2-ChR2-GFP was injected into the VIS to enable ChR2-GFP expression in cells projecting to VIS including ACAvis projection neurons. Averaged firing activity of ChR2 (-) ACA cells in adolescent and adult animals (light blue, n=448 cells from 5 adolescent biologically independent mice, gray, n= 347 cells from 3 adult biologically independent mice, Two-tailed unpaired t-test, $t_{793}$=8.768, ****$P$=0.100x$10^{-14}$). Horizontal lines within violin plots represent median values. Source data are provided as a Source Data file.

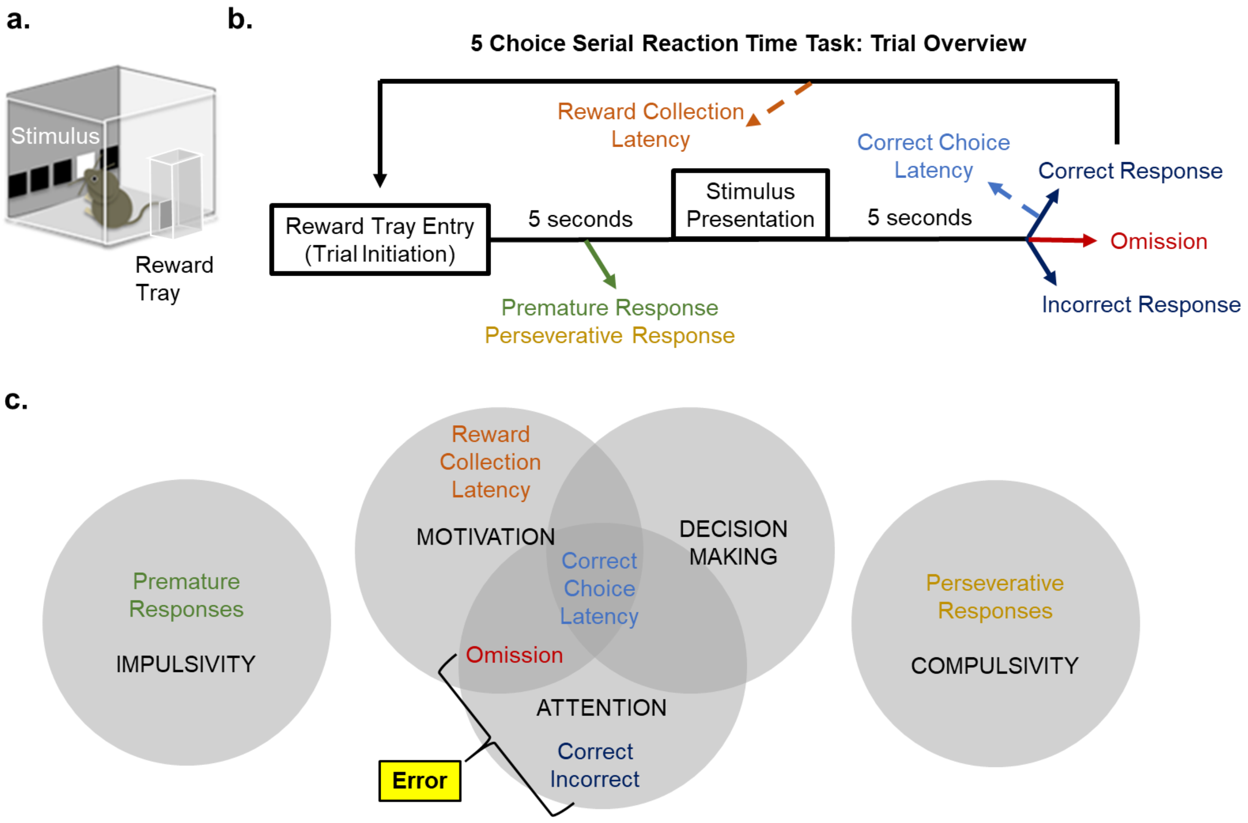


**Supplementary Figure 4. Experimental overview of the 5-Choice Serial Reaction Time Task (5CSRTT) assay of attention (related to Figure 5): a.** Task chambers are equipped with 5 touch-screens, of which one will present as the stimulus when it is lit. The mouse receives a liquid reward from the reward tray when the stimulus is nose-poked. **b**. Trial overview. The mouse initiates a trial by reward tray entry. Multiple performance metrics are gathered automatically through nose pokes and infrared beam breaks throughout the duration of a trial. **c.** Metrics are holistically evaluated to assay overlapping cognitive functions that contribute to multiple aspects of performance with “Error” as the main readout of attentional capacity.

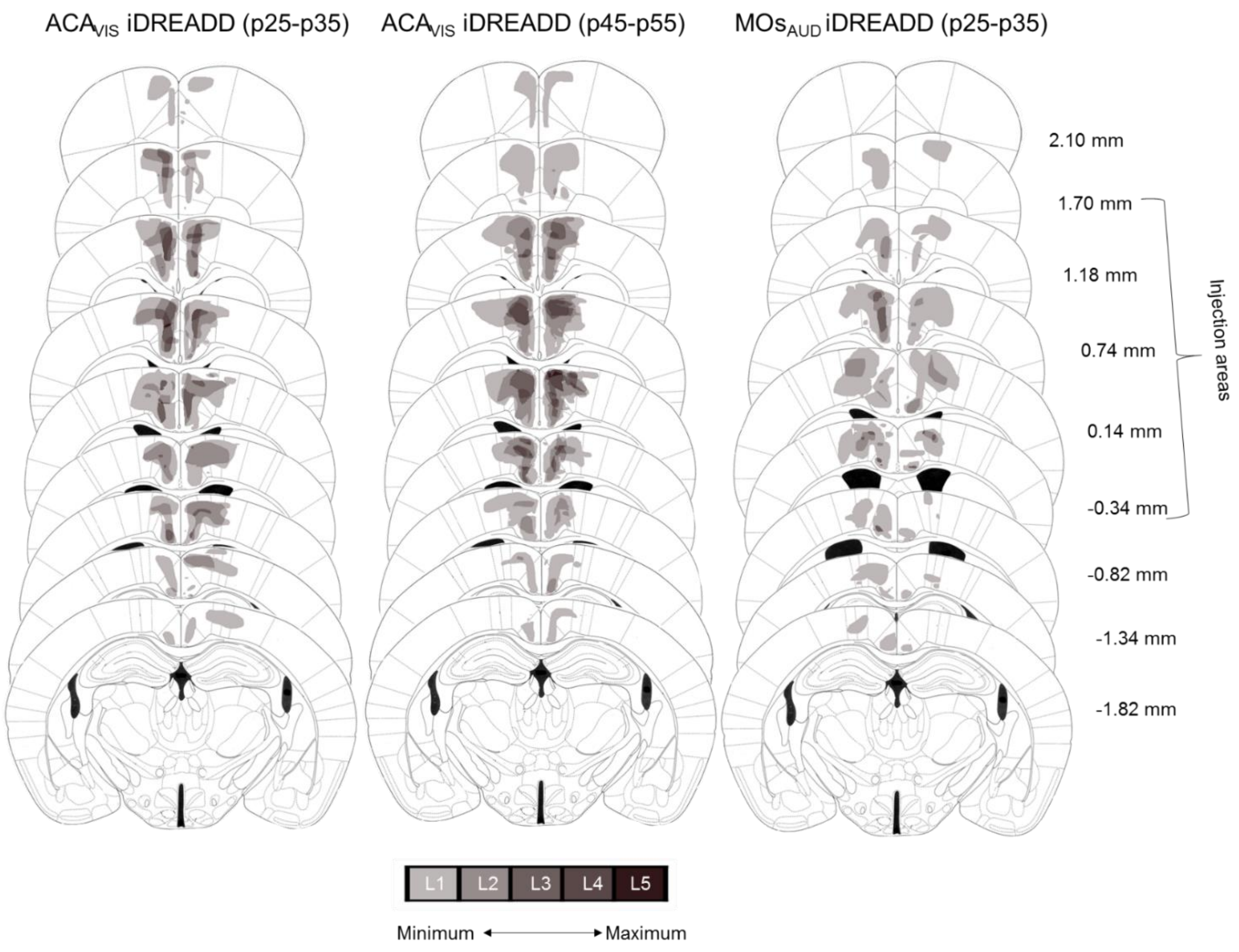


**Supplementary Figure 5. Viral spread validation of iDREADD expression in frontal top-down projection neurons in mice that underwent behavioral testing (related to Figure 5).** Viral spread across the frontal cortex showing expression primarily in expected frontal cortex. To analyze the anterior-posterior spread of virally infected cells, signal intensity was analyzed across nine specific Bregma areas including injection areas indicated with asterisks. All groups were assigned five quartiles representing increasing levels of signal intensity. Signal intensity was determined by the quartile and the number of mice with overlapping viral expression in a given area.

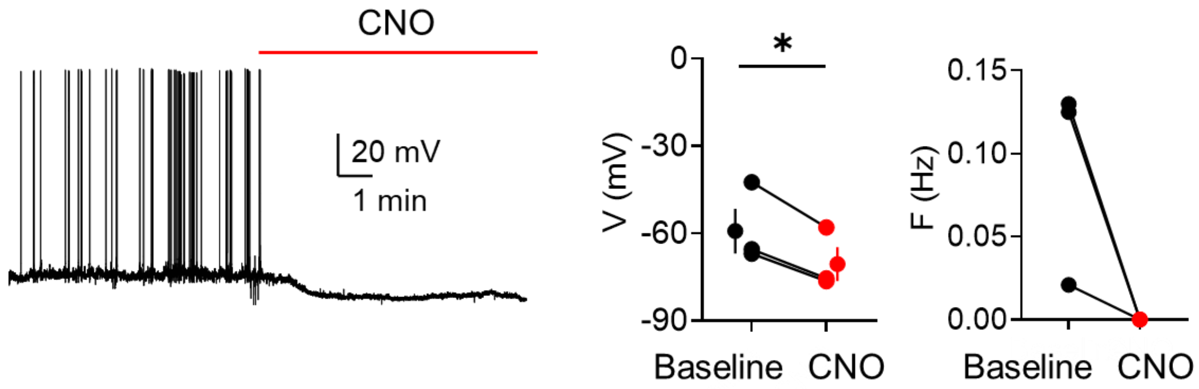


**Supplementary Figure 6**. **Electrophysiologic validation of inhibitory DREADD-mediated top-down inhibition (related to Figure 5)**. (left) Representative whole-cell recording from an adolescent $ACA_{VIS}$ top-down projection neuron in ACA slice upon CNO bath application at p25 following p12 viral injection of inhibitory DREADD. (middle) Membrane potential before (baseline) and during CNO application (CNO) (two-tailed paired *t*-test, $t_2$=6.070, *$P$=0.0261, n=3 cells from 3 biologically independent mice). Markers with error bars represent mean value +/- s.e.m (average age p31.7 ± 0.9). (right) Spontaneous spike frequency of top-down $ACA_{VIS}$ neurons before and during CNO perfusion *in vitro*. Source data are provided as a Source Data file.

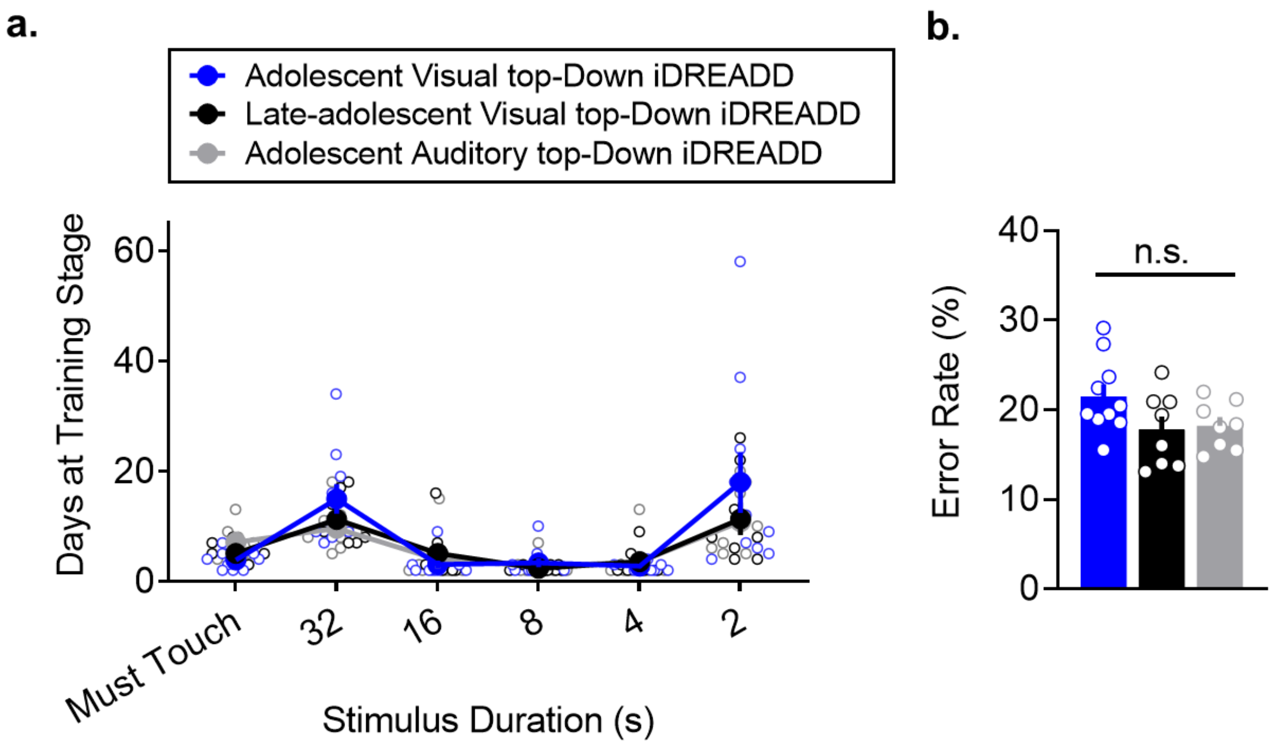


**Supplementary Figure 7. Chemogenetic suppression of frontal top-down projection neurons does not impact 5CSRTT acquisition or baseline training performance (related to Figure 5). a.** iDREADD does not lead to changes in acquisition of the 5CSRTT (Two-way RM ANOVA, effect of group, $F_{2,23}$=0.714, $P$=0.5000, n=10,8,8 biologically independent mice). **b.** iDREADD does not produce differences in baseline performance in rate of combined accuracy and omission errors at 2-s stimulus duration prior to testing (one-way ANOVA $F_{2,23}$=2.708, $P$=0.0879, n=10,8,8 biologically independent mice). Data are presented as mean +/- s.e.m. Source data are provided as a Source Data file.

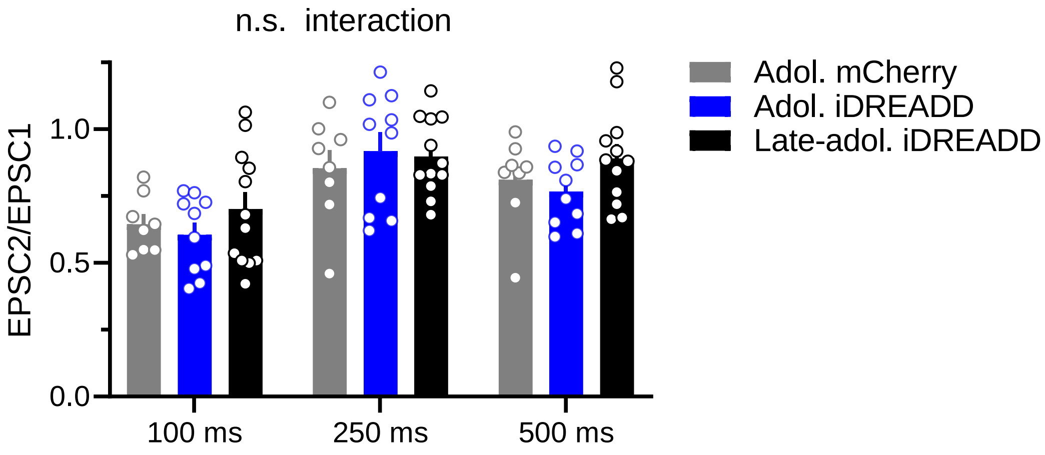


**Supplementary Figure 8: Chemogenetic suppression of frontal top-down $ACA_{VIS}$ projection neuron activity during adolescence does not impact short-term dynamics of local synaptic inputs onto $ACA_{VIS}$ neurons (related to Figure 6).** Two-way RM ANOVA, time x group interaction: $F_{4,54}$=1.611, $P$=0.185, n=10 cells from 5 biologically independent mice for adolescent iDREADD (blue bars); n=8 cells from 3 biologically independent mice for adolescent mCherry (gray bars), and n=12 cells from 6 biologically independent mice for post-adolescent iDREADD (black bars). Data are presented as mean +/- s.e.m. Source data are provided as a Source Data file.

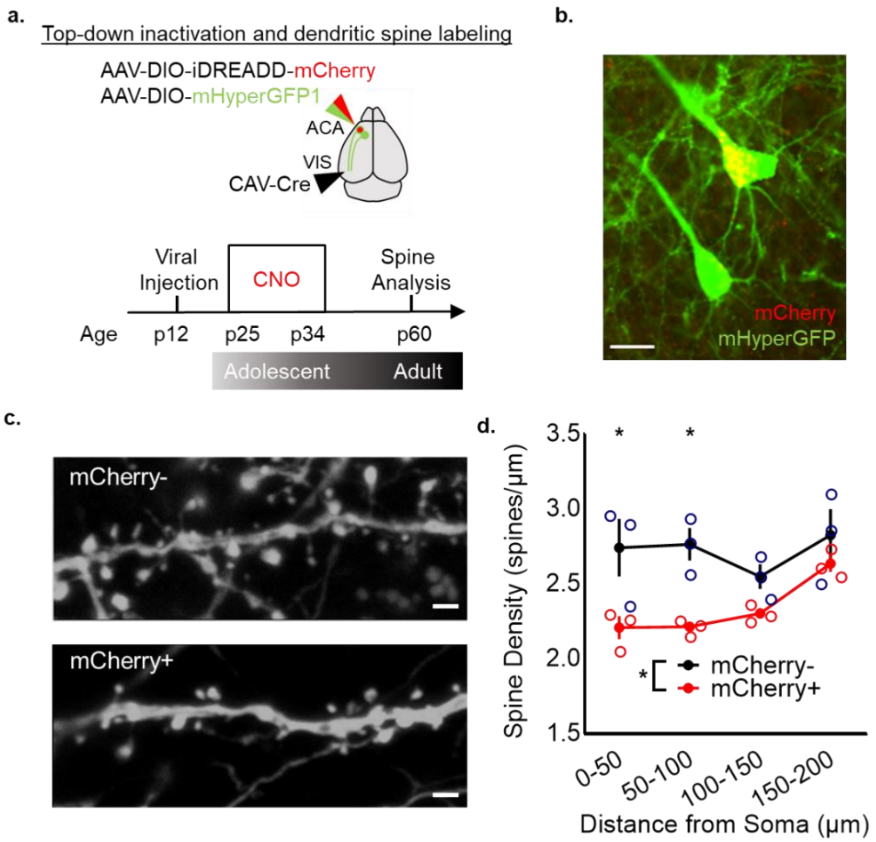


**Supplementary Figure 9. Chemogenetic suppression of $ACA_{VIS}$ projection neurons during adolescence results in excessive loss of dendritic spines in adults (related to Figure 6). a.** Schematic of injection strategy to introduce eGFP selectively into $ACA_{VIS}$ projections and inhibitory DREADD (iDREADD) in only part of these projection neurons for each animal. Viruses were injected at p12 to allow adolescent expression of iDREADD. Mice were injected 10 mg/kg CNO in normal saline twice a day from p25 to p34 and dendritic spines were imaged at adulthood. Spines were analyzed based on their distance from the soma and binned every 50 μm. 4 dendrites/neuron from two iDREADD-expressing neurons (mCherry+) and two mCherry- neurons per mouse were imaged and analyzed for within animal differences. **b.** Representative image of mCherry+ and mCherry- fluorescently labeled top-down neurons (scale bar = 20 μm). Experimental images were obtained from 3 mice with similar results obtained. **c.** Proximal dendrites within the same animal (scale bar = 1μm). Experimental images were obtained from 3 mice with similar results obtained. **d.** Proximal dendrites on mCherry+ neurons show a reduction in total spine density (two-way RM ANOVA, effect of group, $F_{1,4}$=9.966, $P$=0.0343, post-hoc analysis Sidak's multiple comparison test, 0-50 μm distance: *$P$=0.0139, 50-100 μm distance: *$P$=0.0111, 100-150 μm distance: $P$=0.441, 150-200 μm distance: $P$=0.6658 n=3, 3 biologically independent mice). Data in d are presented as mean +/- s.e.m. Source data are provided as a Source Data file.

| | | | |
|---|---|---|---|
| CTXpl | Cortical Plate | | |
| | ISO | Isocortex | |
| | | FRP | Frontal Pole |
| | | AI | Insular Cortex |
| | | ORB | Orbitofrontal Cortex |
| | | ILA | Infralimbic Cortex |
| | | PL | Prelimbic Cortex |
| | *Local* | MOs | Secondary Motor Cortex |
| | *Local* | ACA | Anterior Cingulate Cortex |
| | | MOp | Primary Motor Cortex |
| | | SS | Somatosensory Cortex |
| | | GU | Gustatory Area |
| | | RSP | Retrosplenal Cortex |
| | | VISC | Visceral Area |
| | | AUD | Auditory Cortex |
| | | TEa | Temporal Cortex |
| | | ECT | Entorhinnal Cortex |
| | | PERI | Perirhinal Cortex |
| | | PTLp | Posterior Parietal Cortex |
| | | VIS | Visual Cortex |
| | OLF | Olfactory Areas | |
| | HIP | Hippocampal Formation | |
| BS/IB | Brain Stem | | |
| | Interbrain (diencephalon) | | |
| | | HYP | Hypothalamus |
| | | TH | Thalamus |
| | | DOR | Dorsal Thalamus |
| | | EPI | Epithalamus |
| | | RT | Thalamic Reticular Nucleus |
| | | GENv | Ventral Thalamus |
| | MB | Midbrain | |
| | HB | Hindbrain | |
| CNU | Cerebral Nuclei | | |
| | STR | Striatum | |
| | PAL | Pallidum | |
| CTXsp | Cortical Subplate | | |

**Supplementary Table 1. Hierarchical table of abbreviations for brain regions.** Indented regions are subregions of the outdented region above.
